# Temperature vaulting: A method for screening of slow- and tight-binding inhibitors that selectively target kinases in their non-native state

**DOI:** 10.1101/2025.01.15.632643

**Authors:** Sora Suzuki, Koji Umezawa, Gaku Furuie, Masaki Kikuchi, Daichi G. M. Nakamura, Nanae Fukahori, Ninako Kimura, Masato Yamakawa, Takashi Niwa, Takashi Umehara, Takamitsu Hosoya, Isao Kii

## Abstract

A polypeptide folds into its protein tertiary structure in the native state through a folding intermediate in the non-native state. The transition between these states is thermodynamically driven. A folding intermediate of dual-specificity tyrosine phosphorylation-regulated kinase 1A (DYRK1A) autophosphorylates intramolecularly, whereas DYRK1A in the native state no longer catalyzes this reaction. The alteration in substrate specificity suggests a conformational transition of DYRK1A during its folding process. Consistent with this hypothesis, we identified FINDY (**1**), which inhibits the intramolecular autophosphorylation but not the intermolecular phosphorylation, suggesting that the folding intermediate possesses an alternative inhibitor-binding site. Meanwhile, it remains an issue that the methods for approaching the alternative binding site require an intricate assay tailored to the individual target. Here we show a method, designated as “temperature vaulting,” for screening the non-native-state-targeted inhibitors of DYRK1A. Transient heating of recombinant DYRK1A protein drove the reversible transition between the native state and the non-native state targeted by FINDY (**1**). At physiological temperature, FINDY (**1**) slowly bound to the DYRK1A protein. These results indicate that transient heating accelerates the slow-binding process by assisting the protein to overcome the high-energy barrier leading to the target non-native state. The energy barrier also slowed down the dissociation process, resulting in tight binding between DYRK1A and FINDY (**1**). Furthermore, this study suggests that the dissociation rate underlies the inhibition selectivity of FINDY (**1**) between DYRK1A and its family kinase DYRK1B. This method enables the identification of slow- and tight-binding inhibitors that have been missed in conventional assays.

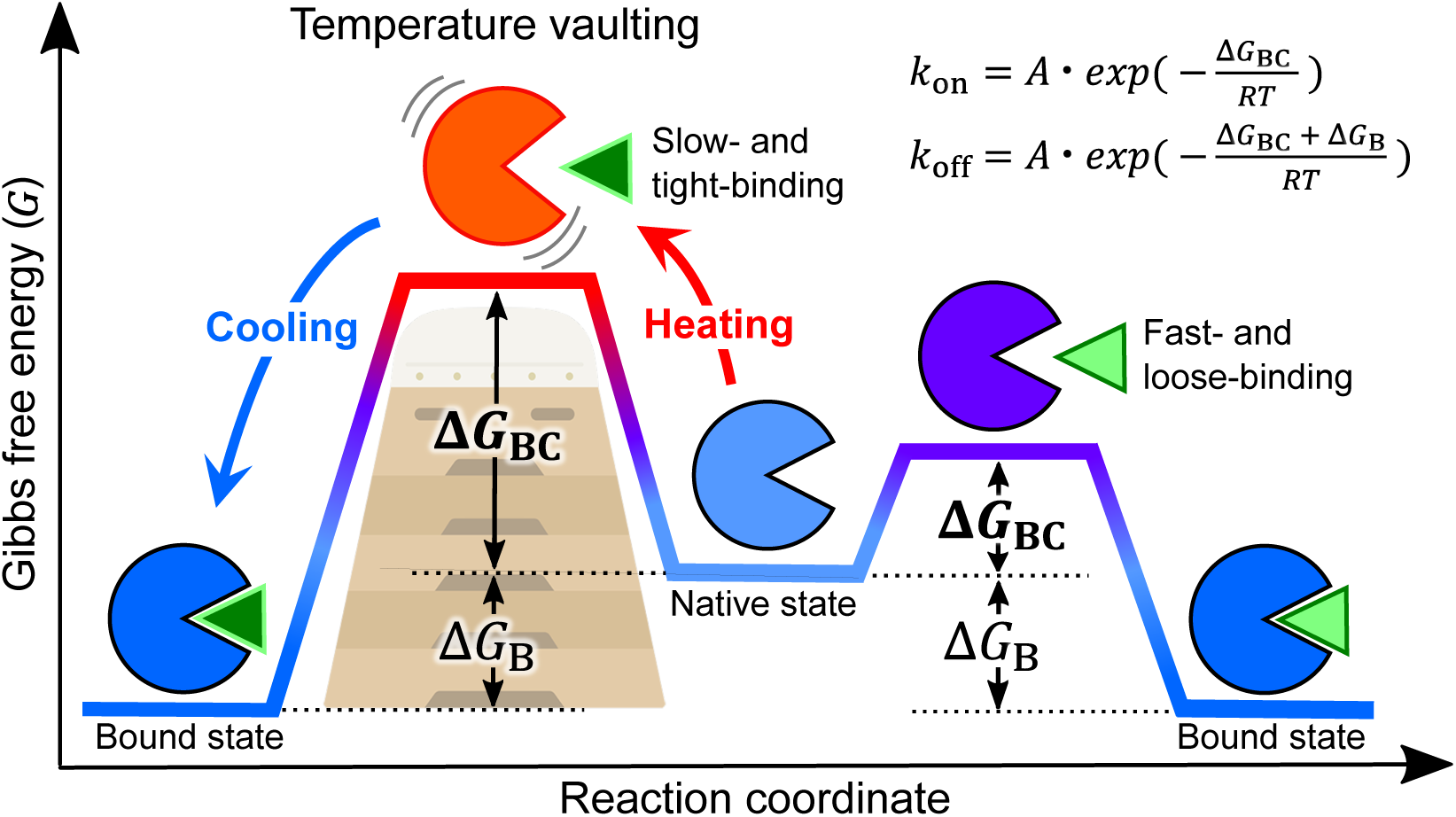

**HIGHLIGHTS:** A temperature vaulting method facilitated the screening of a slow- and tight-binding inhibitor targeting DYRK1A in the non-native state.

A structure–activity relationship study of FINDY (**1**) demonstrated that a slight alteration in chemical structure impacts the inhibition kinetics.

A molecular dynamics simulation of DYRK1A revealed binding-ready conformations for FINDY (**1**).

The dissociation rate may underlie the inhibition selectivity of FINDY (**1**) between DYRK1A and DYRK1B in cellular systems.

## INTRODUCTION

A polypeptide folds into a protein with a particular conformation to execute its biological functions. According to the folding funnel hypothesis,^1^ the folding pathway is represented by the transition from the random coil state of a polypeptide at the edge of the funnel to the bottom, where the native state with its free energy minimum exists (Figure 1A). Along this pathway, a protein also adopts a folding intermediate, which can exist for a while in the non-native local minima within the funnel.^1^ The folding pathway is not unidirectional to the native state, but rather bidirectional. The potential of a protein conformation in each state is given as a function of the Gibbs free energy, expressed in terms of enthalpy, entropy, and temperature.^2–4^ Thus, the ambient temperature governs the proportion of each state within the funnel.^5–7^

**Figure 1.**
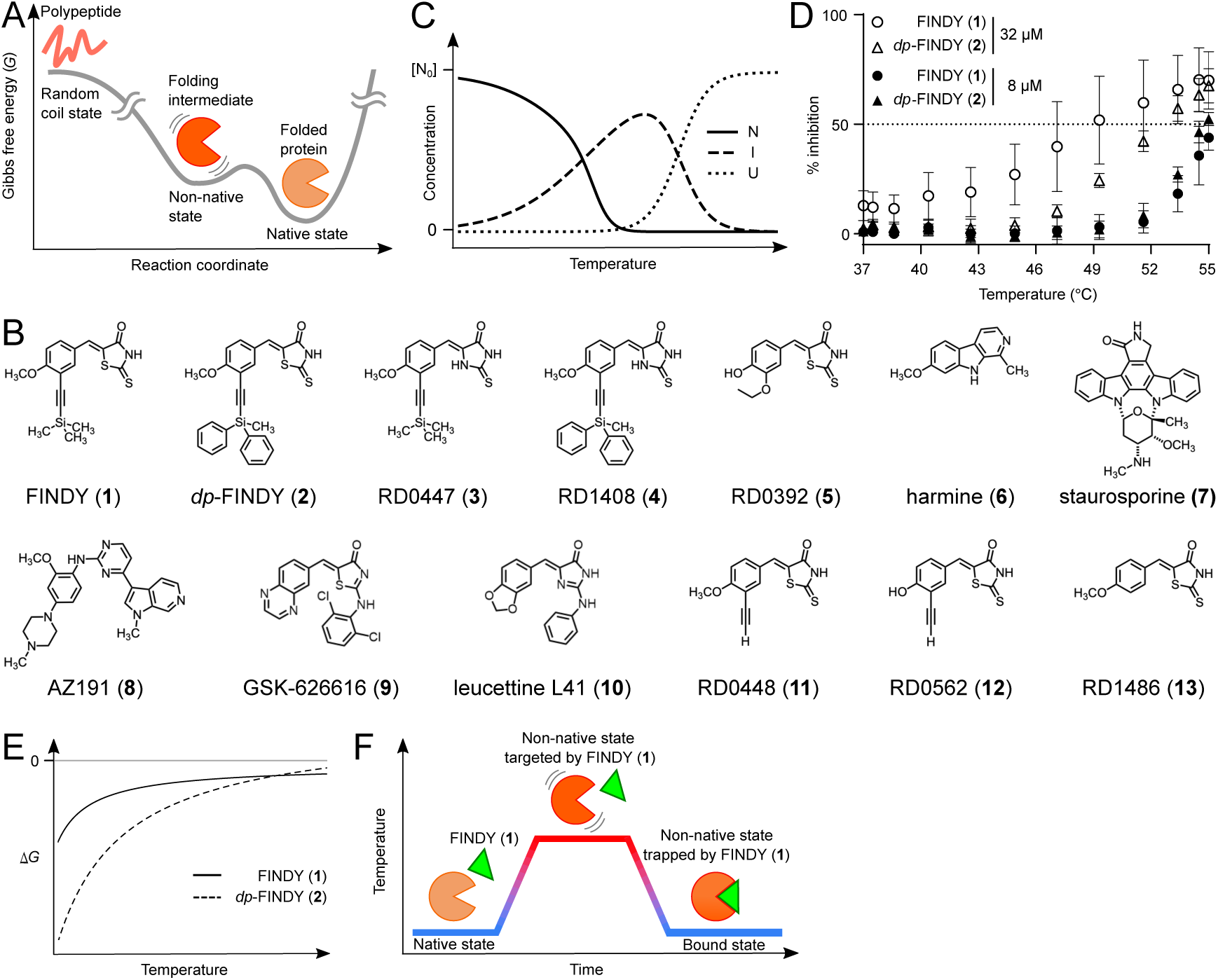
Concept of the protein kinase DYRK1A in the heat-evoked non-native state targeted by FINDY (**1**). (A) Free energy landscape of kinase domain folding. (B) Chemical structures of the compounds used in this study. (C) The concentrations of native state (N), intermediate state (I), and unfolded polypeptide (U) are plotted against temperature according to eqs 9, 10, and 11, respectively. [N_0_] is the sum of [N], [I], and [U]. (D) Inhibition of the DYRK1A protein catalyzed substrate phosphorylation by either FINDY (**1**) or *dp*-FINDY (**2**). The data are presented as mean ± S.D. of three independent experiments measured in duplicate. (E) The differences in Gibbs free energy (Δ*G*) between the non-native state and either the complex with FINDY (**1**) or *dp*-FINDY (**2**) are plotted against temperature according to eq 13. (F) A vaulting box scheme of the transient heating of the DYRK1A protein (circle with a pocket) in the presence of FINDY (**1**). FINDY (**1**) does not bind to the DYRK1A protein in the native state. At a high temperature, the DYRK1A protein undergoes thermodynamic fluctuation, exposing the hidden binding site. The subsequent cooling step stabilizes the complex with FINDY (**1**). The transiently heated protein mixture was subjected to the conventional kinase assay.

To disrupt folding or correct misfolding of disease-related proteins, folding intermediates in the non-native state have been considered druggable as alternatives to folded proteins in the native state.^8–11^ As a protein ascends the layers of the funnel from the bottom to the top, the number of possible conformational states increases, thereby enhancing its conformational flexibility. Conformational flexibility in the inhibitor-binding site has been demonstrated to prolong the residence time (1/*k*_off_, where *k*_off_ is the rate constant for dissociation of the inhibitor–target complex).^12,13^ A prolonged residence time significantly impacts the drug potency and dosing frequency.^12,14–18^

Protein kinases, important pharmaceutical targets, adopt a number of conformational states with distinct catalytic and binding activities.^19–22^ Some of the kinases catalyze intramolecular autophosphorylation on their own residues in their regulatory element.^23–25^ DYRK1A and GSK3β have been demonstrated to autophosphorylate intramolecularly. These autophosphorylations are catalyzed by a folding intermediate in the non-native state but not by the kinases in their native states.^26–28^ In addition, the autophosphorylation-prone intermediate of DYRK1A displays different sensitivity to small-molecule inhibitors compared to its native state,^27^ suggesting conformational flexibility in the inhibitor-binding site. Focusing on this cryptic site, we identified FINDY (**1**) that selectively inhibits the intramolecular autophosphorylation of DYRK1A but not the intermolecular substrate phosphorylation (Figure 1B).^28^ Collectively, these studies demonstrated the existence of a catalytically-active folding intermediate with a conformation distinct from the protein in the native state.

To identify FINDY (**1**) in the previous study, we developed a screening method using living cells with inducible expression of DYRK1A.^28^ This method, however, was not practical for application to other proteins because of its low throughput. Other possible approaches include biophysical measurement of a target protein, which typically requires an expensive dedicated instrument and substantial quantities of purified protein.^29–32^ These issues descending from the above methodologies have become a bottleneck in drug discovery targeting the non-native state.

To alleviate this bottleneck, we established a method based on thermodynamic equilibrium of protein folding states, which could leverage conventional assays widely used in drug screening without reducing robustness. This method adopted temperature control that have been utilized to easily control the equilibrium. A simple kinase assay with this method allowed for quantitative structure–activity relationship analysis of folding-intermediate-selective inhibitors of DYRK1A by using a recombinant DYRK1A kinase domain. This method also accelerated binding during the second step of a two-step slow-binding inhibitor. Furthermore, we found that inhibitors whose inhibitory potency was enhanced by this method bound tightly to the DYRK1A protein. Thus, this method enables the rapid identification of slow- and tight-binding inhibitors targeting kinases in their non-native state.

## RESULTS

### Thermodynamics of DYRK1A in the non-native state targeted by FINDY (1)

We assumed that the DYRK1A folding intermediate targeted by FINDY (**1**) exists in the non-native state, represented as follows:

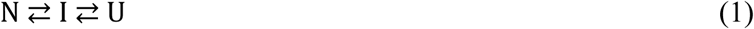

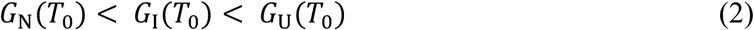

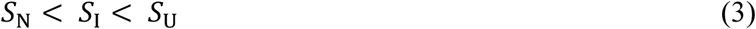

where N, I, and U are the folded protein in the native state, the folding intermediate in the non-native state, and the unfolded polypeptide, respectively. The thermodynamic potential of a protein conformation is given as the Gibbs free energy *G* of each state.^2,7^ *T* indicates the absolute temperature, and *T*_0_ denotes the temperature at which the population of N reaches its maximum. *S* is the entropy of each state. The difference in the Gibbs free energy Δ*G* between any two states determines the thermodynamic potential of the protein conformation in each state under constant pressure, represented as eq 4:

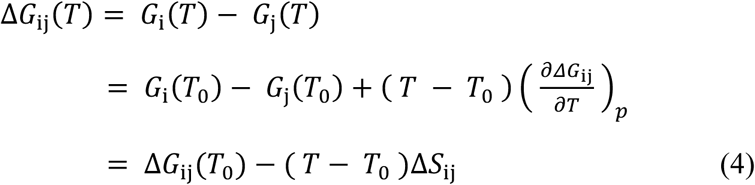

where i and j denote either N, I, or U. The difference in the entropy Δ*S*_ij_ is formulated as a function of *T* and Δ*Q*_ij_, the difference in the heat capacity (eq 5), and substituted into eq 4 to give eq 6.

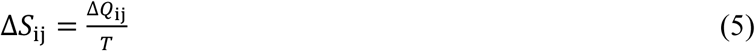

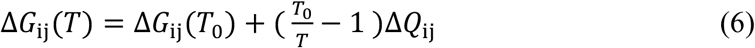

Δ*G*_ij_(*T*) is also represented as follows using the equilibrium constant 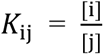:

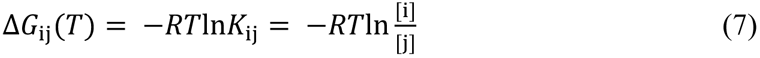

where *R* denotes the gas constant. The concentrations of N, I, and U, denoted as [N], [I], and [U], respectively, are represented as follows:

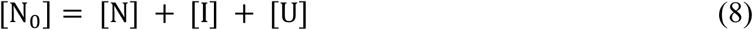

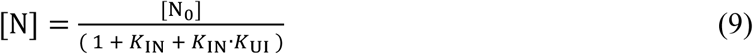

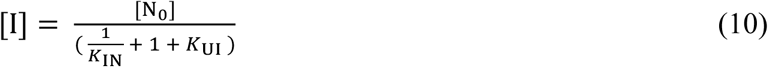

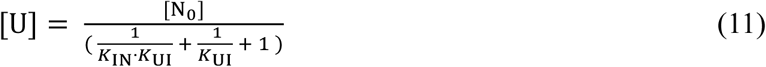

where [N_0_] is the total concentration of the DYRK1A protein. The concentrations plotted against *T* suggest a temperature at which [I] reaches a maximum (Figure 1C). To validate this theory experimentally, we examined whether FINDY (**1**) and *dp*-FINDY (**2**), which are selective inhibitors of the folding intermediate of DYRK1A (Figure 1B),^28,33^ inhibited the substrate phosphorylation activity of recombinant DYRK1A protein at high temperatures. These compounds inhibited it in a temperature-dependent manner (Figure 1D). No difference in the inhibitory potency at 8 μM between FINDY (**1**) and *dp*-FINDY (**2**) was detected; however, FINDY (**1**) was more potent than *dp*-FINDY (**2**) at 32 μM (Figure 1D). These results are inconsistent with that *dp*-FINDY (**2**) showed 66-fold stronger inhibitory potency than FINDY (**1**) in the cell-free autophosphorylation assay.^33^

To understand this inconsistency, we estimated the temperature dependency of the inhibitor-binding equilibrium, using eq 12.

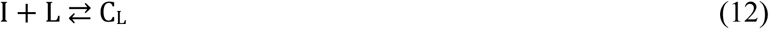

where L denotes either FINDY (**1**) or *dp*-FINDY (**2**), and C_L_ indicates the complex of the intermediate bound with L. The difference in the Gibbs free energy Δ*G* between C_L_ and I is represented using eq 6 as follows:

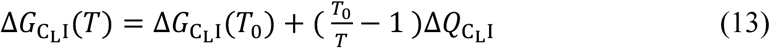

The temperature dependency of the difference in the inhibitory potency between FINDY (**1**) and *dp*-FINDY (**2**) is represented as the difference between Δ*G*_*C*_*dp*-FINDY_1_(*T*) and Δ*G*_*C*_FINDY_1_(*T*) (eq 14):

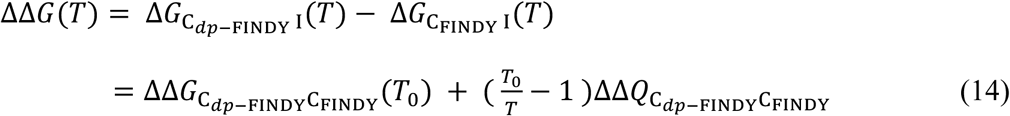

ΔΔ*G*_*C*_*dp*-FINDY_*C*_FINDY__ (*T*_0_) is less than zero because *dp*-FINDY (**2**) inhibited the DYRK1A protein more strongly than FINDY (**1**).^33^ If ΔΔ*Q*_*C*_*dp*-FINDY_*C*_FINDY__ is greater than zero, ΔΔ*G*(*T*) decreases as *T* rises, which indicates that *dp*-FINDY (**2**) becomes more potent than FINDY (**1**) at higher temperatures, being in conflict with the experimental result in Figure 1D. Thus, ΔΔ*Q_Cdp_*_-.FINDY_*_C_*_FINDY_ must be less than zero. A graphical visualization of eq 13 for either FINDY (**1**) or *dp*-FINDY (**2**) is shown in Figure 1E, in which a negative value of ΔΔ*G*(*T*) increases and turns to a positive as *T* rises. Thus, the inhibitory potency of the inhibitors at high temperatures does not correspond to that at physiological temperature.

### Temperature vaulting method targeting the non-native state

To detect the physiological inhibitory potency, we developed a “temperature vaulting” method. In this method, the mixture containing the DYRK1A protein and the selective inhibitor is transiently heated (Figure 1F). The heating time, which should be minimized because prolonged heating causes misfolding and aggregation, was set to allow sufficient for the mixture in the 96-well PCR format to reach the set temperature (Figure S1). In this study, the heating time was set to 20 sec. The heating and cooling rates were set to the maximum of the thermal cycler at measured rates of 4.9 ℃·sec^−1^ and 4.7 ℃·sec^−1^, respectively (Figure S2).

To find the temperature that enables the reversible transition, we determined at what temperature the DYRK1A protein was irreversibly misfolded either during transient heating for 20 sec or during continuous heating for 60 min. The kinase activity was lost at 68 ℃ during transient heating and at 58 ℃ during continuous heating (Figure 2A). Because the optimum point should be between these temperatures, we tentatively decided on a heating temperature of 62 ℃, which is approximately the midpoint. The kinase assay with and without transient heating yielded screening window coefficients (Z′-factor) of 0.87 and 0.86, respectively (Figure S3), indicating that transient heating did not affect the robustness of the assay.

**Figure 2.**
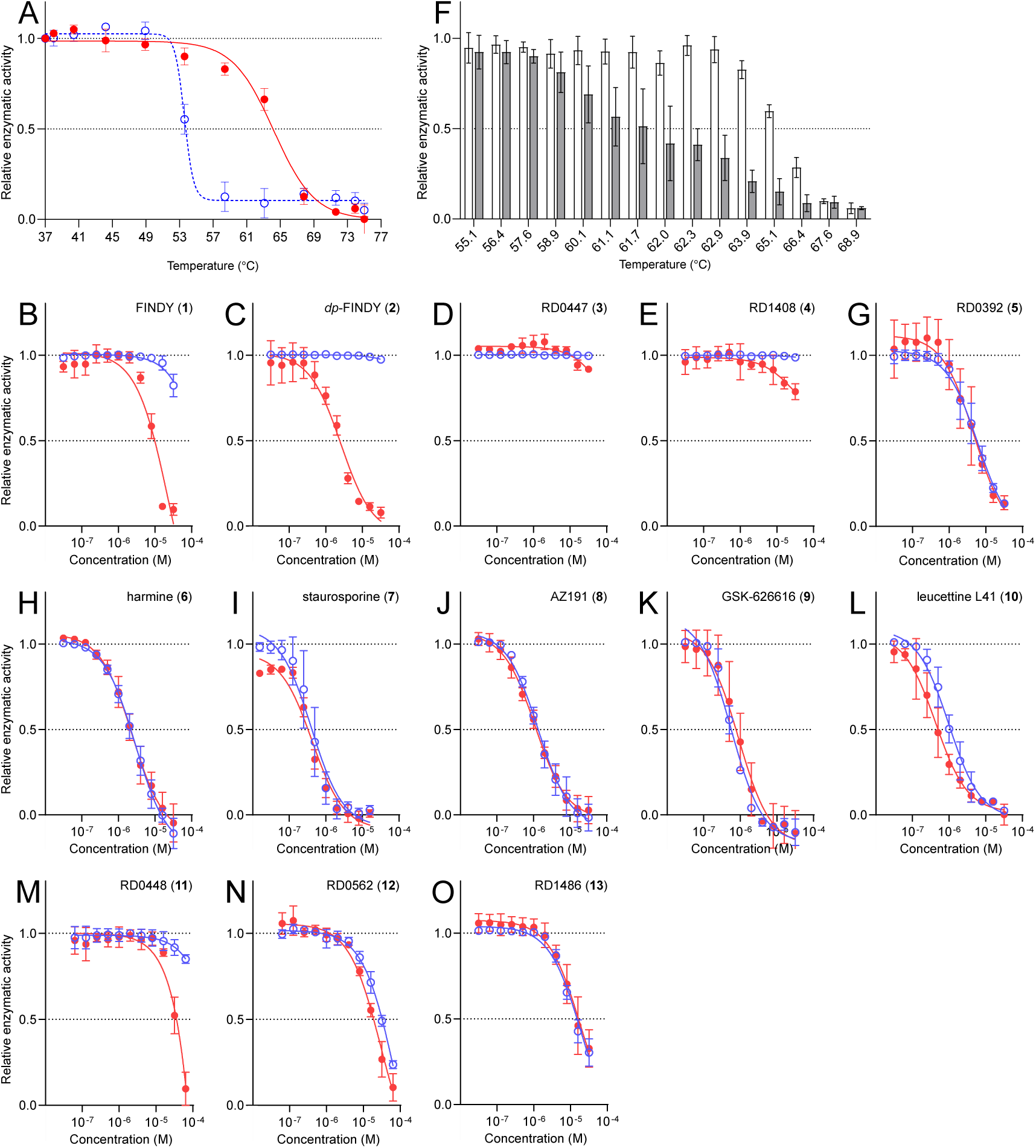
Selective inhibition of the DYRK1A protein using the temperature vaulting method. (A) The closed circles indicate the kinase activities of the DYRK1A protein transiently heated for 20 sec at the indicated temperatures relative to that at 37 ℃. The open circles indicate those continuously heated at the indicated temperatures for 60 min. When the relative activity fell below 0.2, we set this point as the threshold to indicate that the DYRK1A protein became inactive. Representative dose–response curves with Hill slopes are shown. The data are presented as mean ± S.D. of three independent experiments measured in duplicate. (B–E, G–O) The relative kinase activities of the DYRK1A protein in the presence of FINDY (**1**) (B), *dp*-FINDY (**2**) (C), RD0447 (**3**) (D), RD1408 (**4**) (E), RD0392 (**5**) (G), harmine (**6**) (H), staurosporine (**7**) (I), AZ191 (**8**) (J), GSK-626616 (**9**) (K), leucettine L41 (**10**) (L), RD0448 (**11**) (M), RD0562 (**12**) (N), and RD1486 (**13**) (O). The red closed and blue open circles indicate the kinase activities with and without transient heating, respectively. Representative dose–response curves with Hill slopes are shown. The data are presented as mean ± S.D. of three independent experiments measured in duplicate. The IC_50_ values are summarized in Table S1. (F) Determination of the optimum heating temperature. The DYRK1A protein was transiently heated at the indicated temperatures for 20 sec in the presence and absence of FINDY (**1**) at 8 μM (closed and open bars, respectively). The kinase activities relative to that at 37 ℃ in the absence of FINDY (**1**) are displayed on the bar graphs. The data are presented as mean ± S.D. of three independent experiments measured in duplicate.

We examined whether the temperature vaulting method could evaluate the non-native-state-targeted inhibitors of DYRK1A. FINDY (**1**) and *dp*-FINDY (**2**) suppressed the kinase activity in a transient-heating-dependent manner with IC_50_ values of 11 μM and 2.6 μM, respectively (Figure 2B,C). This is consistent with that *dp*-FINDY (**2**) is a more potent selective inhibitor than FINDY (**1**).^33^ When RD0447 (**3**) and RD1408 (**4**) were used as negative control derivatives of FINDY (**1**) and *dp*-FINDY (**2**) (Figure 1B), respectively,^28^ they did not suppress it under either condition (Figure 2D,E). The other derivatives of FINDY (**1**), which have previously been shown to selectively inhibit the intramolecular autophosphorylation of DYRK1A,^33^ also suppressed it in a transient-heating-dependent manner (Figure S4). These results indicated that transient heating could evaluate selective inhibition. In addition, we confirmed whether FINDY (**1**) and *dp*-FINDY (**2**) inhibited the protein once denatured and then refolded. These selective inhibitors did not inhibit it (Figure S5), confirming that the temperature vaulting method enabled a reversible transition between the native state and the non-native state targeted by the selective inhibitors.

To ascertain whether the tentative heating temperature was adequate, we measured the extent to which FINDY (**1**) at 8.0 μM suppressed the kinase activity in the temperature range of 55 ℃ to 69 ℃. The kinase activity was suppressed by more than 50% at temperatures above 62 ℃ (Figure 2F). The inhibitory potency could not be measured reproducibly at temperatures above 66 ℃. Thus, the optimum heating temperature should be between 62 ℃ and 66 ℃. Because the tentative temperature of 62 ℃ was in this range, we continued to use this temperature for our next set of experiments.

In contrast to FINDY (**1**) and *dp*-FINDY (**2**), RD0392 (**5**) that inhibits intermolecular substrate phosphorylation by DYRK1A in the native state^28^ suppressed the kinase activity to the same extent regardless of the conditions (Figures 1B and 2G). We also evaluated the other known inhibitors of DYRK1A.^34^ Harmine (**6**), staurosporine (**7**), AZ191 (**8**), and GSK-626616 (**9**) suppressed it equally regardless of the conditions (Figures 1B and 2H–K). On the other hand, leucettine L41 (**10**) suppressed it more strongly under conditions with transient heating than without it (Figures 1B and 2L). The results indicate that transient heating enhances the potency of the inhibitors, but it depends on their chemical structure.

To investigate the structural requirement for selective inhibition during transient heating, we prepared structural derivatives of FINDY (**1**). RD0448 (**11**), in which the trimethylsilyl group of FINDY (**1**) was replaced with hydrogen (Figure 1B), suppressed the kinase activity in a transient-heating-dependent manner (Figure 2M). RD0562 (**12**), in which both the trimethylsilyl and methoxy groups were replaced with hydrogens (Figure 1B), suppressed it regardless of the conditions (Figure 2N). Thus, the methoxy group is required for the selective inhibition. In addition, the terminal alkyne moiety of RD0448 (**11**) was examined. RD1486 (**13**), in which the alkyne moiety of RD0448 (**11**) was replaced with hydrogen (Figure 1B), suppressed it regardless of the conditions (Figure 2O). Taken together, both the methoxy group and the alkyne moiety underlie the selectivity of RD0448 (**11**) toward the DYRK1A protein in the non-native state.

### Conformation of DYRK1A in the non-native state targeted by FINDY (1)

Our previous study solved the crystal structures of the DYRK1A kinase domain complexed with RD0392 (**5**) and RD0448 (**11**).^33^ Superimposition of these native conformations revealed no significant difference in the root-mean-square distance (RMSD) of 0.221 Å (Figure S6A), which could not clearly explain the observed selective inhibition by RD0448 (**11**) during transient heating. However, the similarity between the complete bound forms does not ensure that between the binding processes. We postulated that the selective inhibition by RD0448 (**11**) may be attributed to targeting of a binding-competent state beyond the energy barrier in the energy landscape, denoted as Δ*G*_BC(RD0448)_ in Figure 3A, which is larger than Δ*G*_BC(RD0392)_. To understand the protein conformation in the high-energy binding-competent states targeted by either FINDY (**1**), *dp*-FINDY (**2**), or RD0448 (**11**), we assessed *in silico* docking models of the selective inhibitors to the ATP-binding pocket of the apo-kinase domain of DYRK1A at high temperatures in a 2-μsec all-atom molecular dynamics (MD) simulation. The RMSD with the initial structure indicated that the simulated conformations under 70 ℃ were maintained (RMSD < 2.5 Å), while those at higher temperature became unfolded (Figure S6B). The conformations that exhibited no collision with the docked ligands in the pocket were counted and displayed as a percentage at the indicated temperatures (Figure 3B). At 25 ℃, 40 ℃, and 55 ℃, the percentages of the conformations without collision relative to the total were low but still present (Figure 3B). At 70 ℃, the percentages increased (Figure 3B). On the other hand, the percentages decreased at 85 ℃ and 100 ℃ due to steric hindrance caused by structural collapse of the pocket (Figure 3B). Principal component analysis (PCA) of the simulated conformations showed that the conformational fluctuation expanded in the PCA space as the temperature rose (Figure S6C). The representative image of the conformation without collision at 70 ℃ is shown as a ribbon diagram in Figure 3C,D. Our previous molecular modeling study suggested steric hindrance between FINDY (**1**) and the ATP-binding pocket of DYRK1A, in which Ile165 was in direct conflict with the bulky hydrophobic trimethylsilyl moiety of FINDY (**1**).^33^ In the present simulation, although substantial differences in the overall fold between these conformations were not observed (Figure 3C), the rotational dynamics of Ile165 was enhanced by heating to make space for ligand binding (Figure 3D). Thus, the MD simulation generated binding-ready conformations for the selective inhibitors.

**Figure 3.**
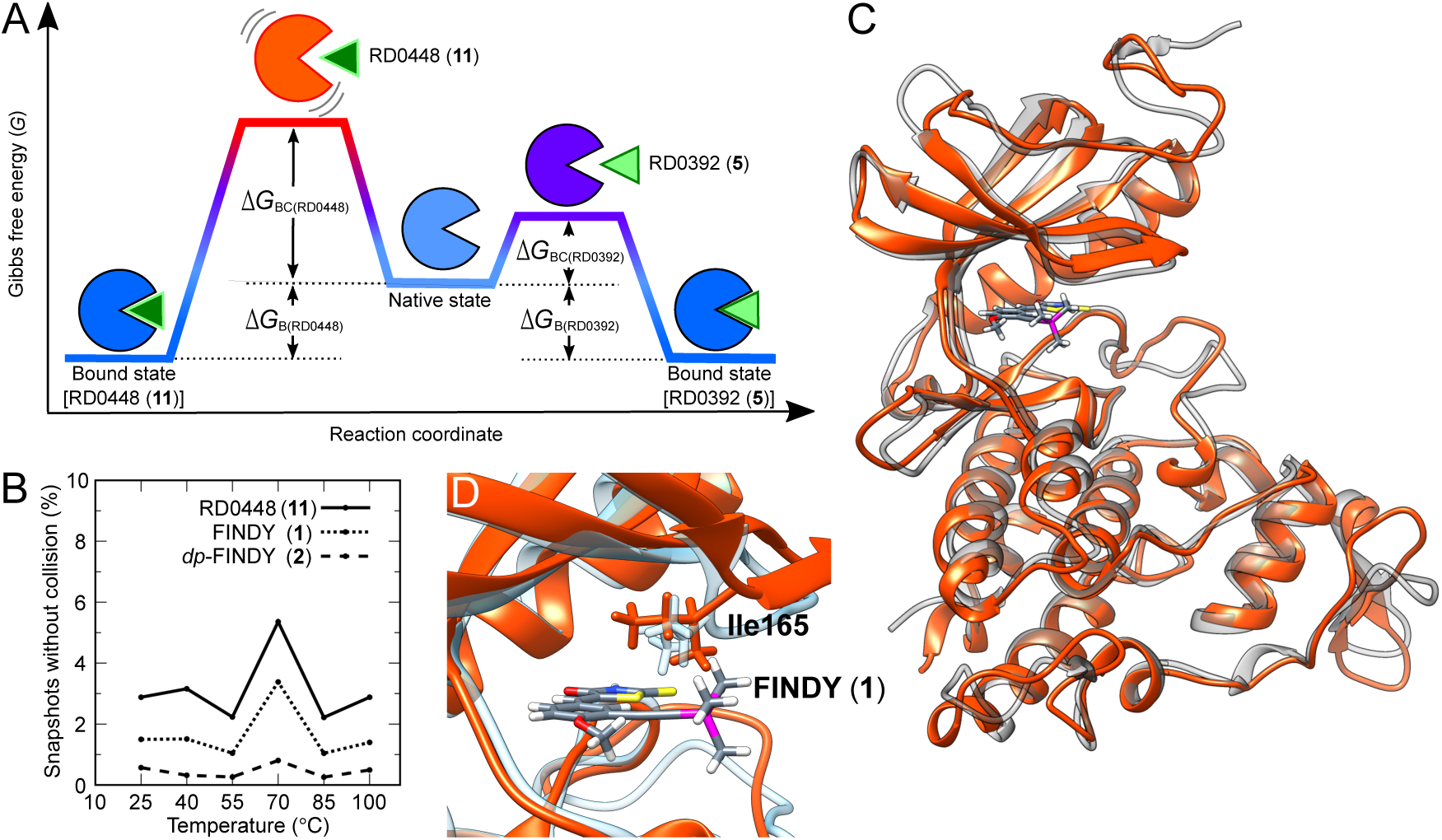
Molecular dynamics simulation of the kinase domain and subsequent molecular modeling for binding to FINDY (**1**). (A) Schematics of the free energy landscape of a one-barrier inhibitor–target binding profile. A one-step model with one free energy barrier is shown. Δ*G*_BC_ indicates the difference in the free energy between the binding-competent state for the inhibitor and the native state. Δ*G*_B_ indicates the difference in the free energy between the native state and the bound state. (B) Appearance probability of the conformations bound with either FINDY (**1**), *dp*-FINDY (**2**), and RD0448 (**11**) without collision in the ensemble simulated at the indicated temperatures. (C) Overlay of the ribbon diagrams of the initial apo form (gray) at 25 ℃ and the representative conformation bound with FINDY (**1**) (orange) at 70 ℃. FINDY (**1**) is shown in stick form. (D) An enlarged view of the ATP-binding pocket in (C). The side chain of Ile165 is shown in stick form.

### Inhibition kinetics of the non-native-state-targeted inhibitors

Although RD0448 (**11**) did not suppress the kinase activity of the DYRK1A protein in the native state (Figure 2M), our previous study demonstrated preparation of the DYRK1A kinase domain complexed with RD0448 (**11**),^33^ indicating that incubation time allowed RD0448 (**11**) to bind to the recombinant protein at the crystallization temperature of 20 ℃. To determine the incubation time required for the binding, mixtures containing the DYRK1A protein and the small molecules were incubated for up to 48 hr at 37 ℃ prior to the kinase reaction. FINDY (**1**) began to suppress the kinase activity as the pre-incubation time increased, and the inhibitory potency reached a plateau at 8 hr (Figure 4A and Table S2). *dp*-FINDY (**2**) and RD0448 (**11**) also inhibited the protein to a greater extent as the time increased (Figure 4B,C and Table S2). RD0447 (**3**) and RD1408 (**4**) did not inhibit it after pre-incubation (Figure 4D,E). On the other hand, RD0392 (**5**) showed the same inhibitory potency over the entire incubation period (Figure 4F and Table S2). Thus, these results suggest that the non-native-state-selective inhibitors exhibit slow-binding kinetics due to the time required for the DYRK1A protein to transition from the native state to the target non-native state. In addition, leucettine L41 (**10**) inhibited the protein at the initial time point of 0 hr with IC_50_ value of 0.74 μM, and the inhibitory potency reached a plateau at 8 hr with IC_50_ value of approximately 0.5 μM (Figure 4G and Table S2). The time-dependent enhancement of inhibitory potency indicates that leucettine L41 (**10**) inhibits DYRK1A by a two-step mechanism with initial loose binding followed by slow constriction to a high-affinity complex. Taken together, transient heating accelerates the slow-binding step of the inhibitors.

**Figure 4.**
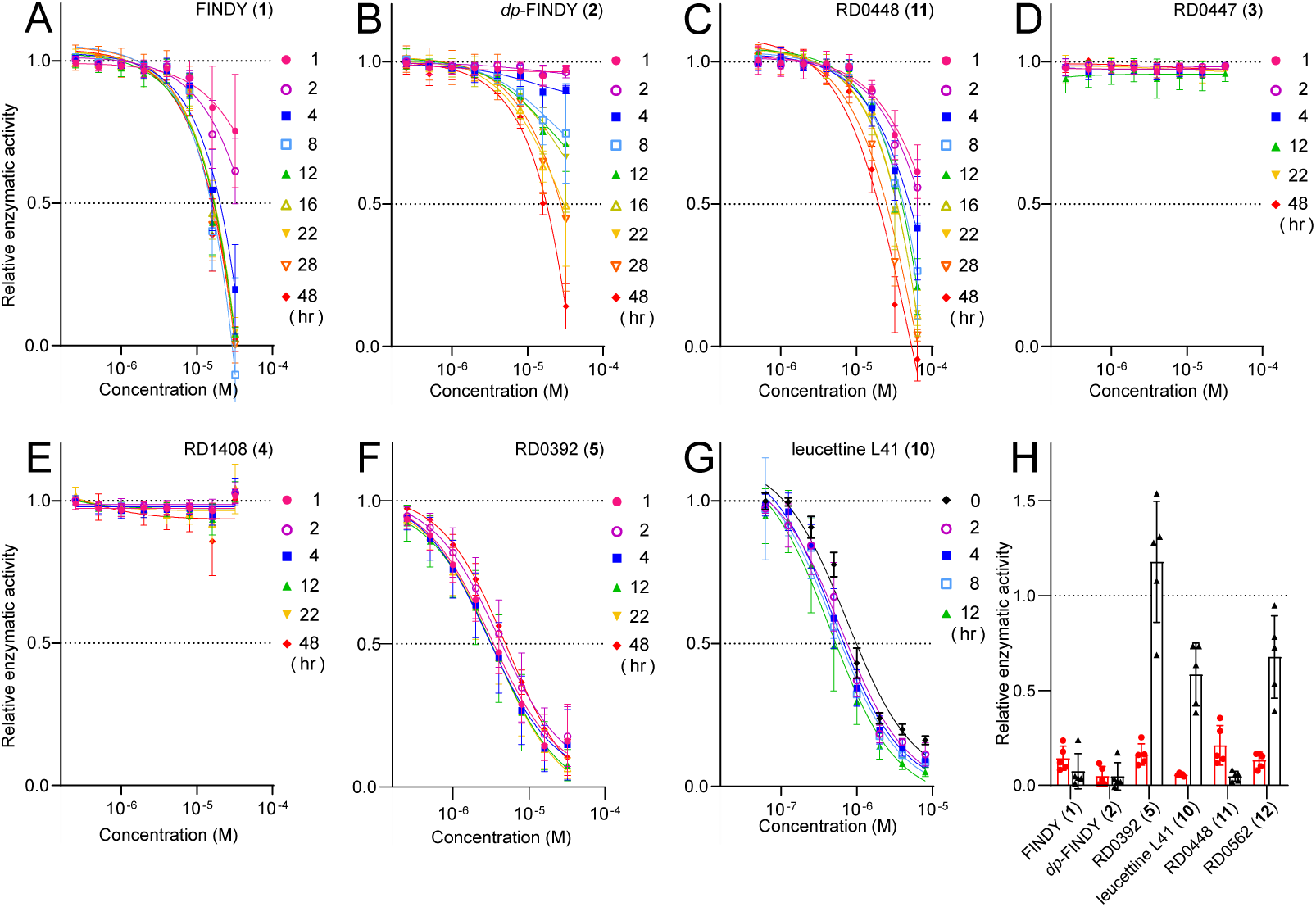
FINDY (**1**) exhibited slow association and dissociation kinetics for the DYRK1A protein. (A–G) The DYRK1A protein with the indicated inhibitors at the indicated concentrations was incubated for the indicated times prior to the kinase reaction. The pre-incubated proteins were reacted with substrate peptide and ATP. The data are presented as mean ± S.D. of three independent experiments measured in duplicate. The IC_50_ values are summarized in Table S2. (H) The DYRK1A protein was transiently heated with either FINDY (**1**), *dp*-FINDY (**2**), RD0392 (**5**), or leucettine L41 (**10**) at 32 μM, and with either RD0448 (**11**) or RD0562 (**12**) at 64 μM, which was then loaded onto a PD-10 desalting column to remove the unbound inhibitors. The red and black open bars indicate the relative kinase activities before and after the gel filtration, respectively. The data are presented as mean ± S.D. of five independent experiments measured in duplicate.

The structural analyses and the slow-binding kinetics of the selective inhibitors suggest that the energy barrier leading up to the non-native binding-competent state cannot be overcome without transient heating or a long pre-incubation step (Figure 3A). It has been demonstrated that destabilization of the binding-competent transition state slows down not only the association rate but also the dissociation rate of the inhibitor–target complex.^12^ According to the energy landscape model shown in Figure 3A, the binding-competent-state destabilization corresponds to an increase in the Δ*G*_BC_, which simultaneously increases the total free energy barrier (Δ*G*_BC_ + Δ*G*_B_) for the dissociation of the complex. In support of this thermodynamics-based model, the slow dissociation of FINDY (**1**) from DYRK1A has been observed in our previous studies.^28,35^ DYRK1A purified from HEK293 cells using a buffer without FINDY (**1**), which were cultured with FINDY (**1**), remained inhibited.^28^ In addition, FINDY (**1**) was detected still bound to the DYRK1A kinase domain purified from *E. coli* cells using a buffer without FINDY (**1**), which were cultured with FINDY (**1**).^35^

To validate the association/dissociation process of the selective inhibitors, the rate constants for association and dissociation of the inhibitors of DYRK1A were calculated. The pre-incubation titration curves in Figure 4A,B and Figure S7A,B were analyzed using a previously reported kinetic model^36,37^ and the parameters were calculated. The association and dissociation rate constants of FINDY (**1**) were 9.4 M^−1^·sec^−1^ and 1.7 × 10^−5^ sec^−1^ at 37 ℃, respectively (Table 1). Those of *dp*-FINDY (**2**) were 2.8 × 10^−1^ M^−1^·sec^−1^ and 2.2 × 10^−7^ sec^−1^ at 37 ℃, respectively (Table 1). We also determined the association and dissociation rate constants of RD0392 (**5**) by surface plasmon resonance (Figure S8), which were calculated to be 2.0 × 10^6^ M^−1^·sec^−1^ and 2.6 × 10^−1^ sec^−1^ at 25 ℃, respectively (Table 1). These results clearly demonstrated that FINDY (**1**) and *dp*-FINDY (**2**) were slow- and tight-binding inhibitors.

**Table 1.** Kinetics and affinities of the inhibitors.

|  | FINDY (1) |  | *dp*-FINDY (2) |  | RD0392 (5) |
| --- | --- | --- | --- | --- | --- |
|  | DYRK1A | DYRK1B | DYRK1A | DYRK1B | DYRK1A |
| $k_{\text{on}}$ ($\text{M}^{-1}\cdot\text{sec}^{-1}$) | 9.4 | $2.6 \times 10^2$ | $2.8 \times 10^{-1}$ | $2.0 \times 10^2$ | $2.0 \times 10^6$ |
| $k_{\text{off}}$ ($\text{sec}^{-1}$) | $1.7 \times 10^{-5}$ | $3.7 \times 10^{-4}$ | $2.2 \times 10^{-7}$ | $1.2 \times 10^{-4}$ | $2.6 \times 10^{-1}$ |
| $K_D$ (M) | $1.8 \times 10^{-6}$ | $1.4 \times 10^{-6}$ | $8.0 \times 10^{-7}$ | $6.0 \times 10^{-7}$ | $1.3 \times 10^{-7}$ |
| Residence time | 17 hr | 45 min | 52 days | 2.4 hr | 3.8 sec |
| $R^2$ | 0.87 | 0.83 | 0.83 | 0.92 | |
| Temperature ($^{\circ}\text{C}$) | 37 | 37 | 37 | 37 | 25 |

$R^2$ : correlation coefficient of the global fitting to eq 15

Since the dissociation rate constants of FINDY (**1**) and *dp*-FINDY (**2**) were too small compared to canonical small-molecule inhibitors, the dissociation rates of the inhibitors were confirmed using a different approach. To easily compare the dissociation rates, we performed a gel filtration experiment to determine whether the inhibitors dissociated from the DYRK1A complex after the unbound inhibitors were removed from the mixture that had been transiently heated. After the removal of the unbound inhibitors by gel filtration, the kinase activity was recovered in the mixture containing either RD0392 (**5**), leucettine L41 (**10**), or RD0562 (**12**) (Figure 4H). The DYRK1A protein transiently heated with RD0392 (**5**) showed greater activity than that heated with solvent after the removal of the inhibitor (Figure 4H), indicating that RD0392 (**5**) was a reversible inhibitor that prevented thermal denaturation of the DYRK1A protein. In contrast, the kinase activity was not recovered in the mixture containing either FINDY (**1**), *dp*-FINDY (**2**), or RD0448 (**11**) (Figure 4H), indicating that these inhibitors tightly bound to the kinase domain. Thus, these results confirmed the slow dissociation of FINDY (**1**) and *dp*-FINDY (**2**) from its complex.

### Destabilization of the binding-competent state slows down the dissociation rate

As illustrated in Figure 3A, slow dissociation could be achieved through binding-competent-state destabilization (the increase in Δ*G*_BC_), bound-state stabilization (the increase in Δ*G*_B_), or a combination of both. Because Δ*G*_B_ quantitatively relates to the dissociation constant *K*_D_ when the state transition reaches equilibrium,^38^ the *K*_D_ values in Table 1 indicate that Δ*G*_B(RD0392)_ was larger than Δ*G*_B(FINDY)_ and Δ*G*_B(*dp*-FINDY)_, suggesting that the bound-state stabilization did not have a significant contribution to the slow dissociation of FINDY (**1**) and *dp*-FINDY (**2**).

To ensure the contribution of Δ*G*_B_, we compared the stability of the complexes, which was governed by their Δ*G*_B_, by measuring the melting temperatures of the DYRK1A kinase domain in the presence of the inhibitors. The kinase domain exhibited melting temperatures of 48.5 ℃ and 56.5 ℃ (Figure S9A), indicating independent thermal denaturation of the N- and C-lobes. Although co-incubation with RD0392 (**5**) gave a positive shift in the melting temperature to a single peak at 59.0 ℃ (Figure S9B), this shift was not observed for FINDY (**1**) and *dp*-FINDY (**2**), where two peaks at 42.9 ℃ and 55.8 ℃ were observed for co-incubation with FINDY (**1**), and a single peak at 57.1 ℃ was observed for *dp*-FINDY (**2**) (Figure S9C,D). The negative control compound RD0447 (**3**) did not affect the melting temperatures, where two peaks at 50.9 ℃ and 57.3 ℃ were observed (Figure S9E). Thus, the slow dissociation appears to be driven by binding-competent-state destabilization rather than bound-state stabilization.

### Inhibition selectivity of FINDY (1) between DYRK1A and DYRK1B

The aforementioned study has provided evidence on the chemical structure required for slow- and tight-binding inhibitors of DYRK1A. The subsequent question in this study was whether slow- and tight-binding inhibition contributed to the selectivity between kinases. We therefore investigated the structure–activity relationship of the inhibitors for inhibition of the recombinant DYRK1B kinase domain, which is highly similar to DYRK1A.^39^ Our previous study demonstrated that FINDY (**1**) rescued the DYRK1A-overexpression-induced developmental malformation of *Xenopus* embryos but did not the DYRK1B-overexpression-induced ones,^28^ indicating that FINDY (**1**) selectively inhibits DYRK1A but not DYRK1B, whereas the mechanism underlying the selectivity has remained elusive.

The heating temperature for the recombinant DYRK1B kinase domain was determined to be 54 ℃ (Figures 5A and S10A). FINDY (**1**), *dp*-FINDY (**2**), and RD0448 (**11**) showed transient-heating-dependent enhancement of the inhibitory potency against DYRK1B (Figure 5B–D), which was the same trend as for DYRK1A (Figure 2B–D). The FINDY derivatives also showed enhancement (Figure S10B). In contrast, leucettine L41 (**10**) did not show heating-dependent enhancement (Figure 5E). The other inhibitors suppressed the kinase activity to the same extent regardless of the conditions (Figure 5F–I and Figure S10C). At the physiological temperature of 37 ℃, FINDY (**1**), *dp*-FINDY (**2**), and RD0448 (**11**) suppressed the kinase activity to a greater extent as the pre-incubation time increased (Figure 5J–L and Table S3). RD0392 (**5**) and leucettine L41 (**10**) showed the same inhibitory potency over the entire incubation period (Figure 5M,N and Table S3). The pre-incubation titration curves in Figures 5J,K and Figure S7C,D were analyzed using the kinetic model and the parameters were calculated. The association and dissociation rate constants of FINDY (**1**) for DYRK1B were 2.6 × 10^2^ M^−1^·sec^−1^ and 3.7 × 10^−4^ sec^−1^ at 37 ℃, respectively (Table 1). Those of *dp*-FINDY (**2**) for DYRK1B were 2.0 × 10^2^ M^−1^·sec^−1^ and 1.2 × 10^−4^ sec^−1^ at 37 ℃, respectively (Table 1). These results indicate that these inhibitors bound to and dissociated from the DYRK1B protein at a faster rate than DYRK1A. A gel filtration experiment demonstrated the correlation between transient-heating-dependent enhancement of inhibitory potency and sustained inhibition after the removal of the unbound inhibitors in DYRK1B as well as DYRK1A (Figure 5O). Taken together, the reason why FINDY (**1**) inhibits DYRK1A but not DYRK1B in the *in vivo* model could be the approximately 22-fold difference in the dissociation rate constants between DYRK1A and DYRK1B.

**Figure 5.**
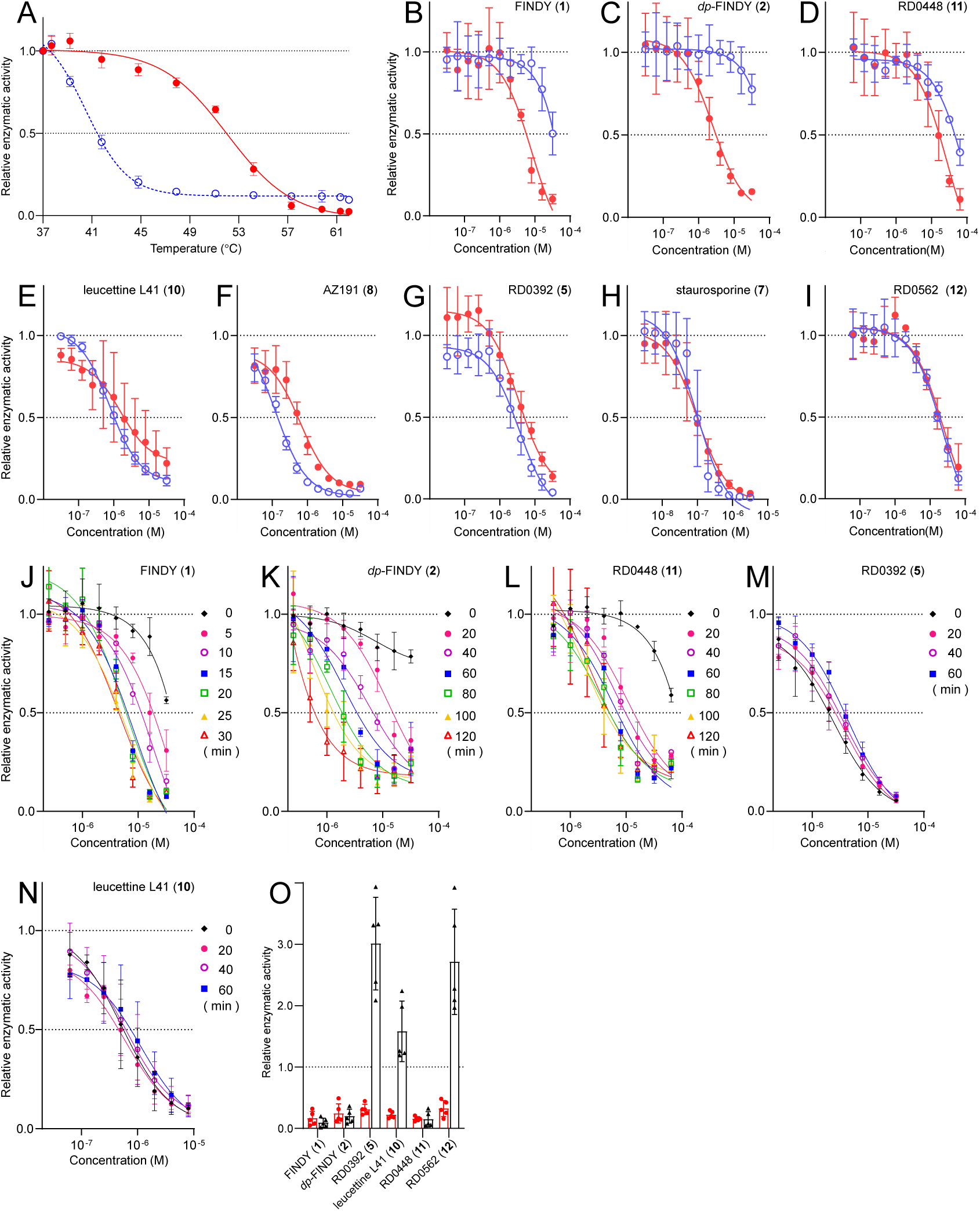
Slow association and dissociation kinetics of FINDY (**1**) for the DYRK1B protein. (A) The closed circles indicate the kinase activities of the DYRK1B protein transiently heated for 20 sec at the indicated temperatures, relative to that at 37 ℃. The open circles indicate those continuously heated at the indicated temperatures for 60 min. When the relative activity fell below 0.2, we set this point as the threshold to indicate that the DYRK1B protein became inactive. Representative dose–response curves with Hill slopes are shown. The data are presented as mean ± S.D. of three independent experiments measured in duplicate. (B–I) The relative kinase activities of the DYRK1B protein in the presence of FINDY (**1**) (B), *dp*-FINDY (**2**) (C), RD0448 (**11**) (D), leucettine L41 (**10**) (E), AZ191 (**8**) (F), RD0392 (**5**) (G), staurosporine (**7**) (H), and RD0562 (**12**) (I). The red closed and blue open circles indicate the kinase activities with and without transient heating, respectively. Representative dose–response curves with Hill slopes are shown. The data are presented as mean ± S.D. of three independent experiments measured in duplicate. The IC_50_ values are summarized in Table S1. (J–N) The DYRK1B protein with the indicated inhibitors at the indicated concentrations was incubated for the indicated times prior to the kinase reaction. The data are presented as mean ± S.D. of three independent experiments measured in duplicate. The IC_50_ values are summarized in Table S3. (O) The DYRK1B protein was transiently heated with either FINDY (**1**), *dp*-FINDY (**2**), RD0392 (**5**), or leucettine L41 (**10**) at 32 μM, and with either RD0448 (**11**) or RD0562 (**12**) at 64 μM, which was then loaded onto a PD-10 desalting column to remove the unbound inhibitors. The red and black open bars indicate the relative kinase activities before and after the gel filtration, respectively. The data are presented as mean ± S.D. of five independent experiments measured in duplicate.

## DISCUSSION

Inspired by thermodynamic equilibrium between protein conformational states, we developed the temperature vaulting method that enables the facile evaluation of inhibitors targeting protein kinases in their non-native state by leveraging the instant kinase assay without affecting its robustness. Inhibitors targeting the non-native state exhibited slow-binding kinetics for DYRK1A in the native state. The slow-binding kinetics was attributed to the time required for the transition from the native state to the non-native state targeted by the selective inhibitors at the physiological temperature of 37 ℃. Thus, the temperature vaulting method allows acceleration of the association of a slow-binding inhibitor to its target protein. This acceleration was observed in the binding of leucettine L41 (**10**). As a reasonable outcome in accordance with the energy landscape model for the binding-competent state, the slow-binding inhibitors exhibited slow dissociation kinetics, forming tight-binding complexes. The high-energy barrier leading up to the binding-competent state, Δ*G*_BC_ in Figure 3A, slows down not only the association process but also the dissociation process.^12^ This study developed a rational strategy for screening slow- and tight-binding inhibitors of DYRK1A.

The MD simulation at high temperature revealed binding-ready conformations in the non-native state. The probability of the conformations became largest at 70 ℃ whereas the overall fold remained similar to the crystal structure in the native state. At lower temperatures, steric hindrance between FINDY (**1**) and Ile165 in the ATP-binding pocket entrance was not resolved. At higher temperatures, steric hindrance was increased by structural collapse of the pocket. Thus, optimization of simulation temperature was necessary to expose the alternative binding site. Heating at 70 ℃ promoted conformational fluctuation including reorientation of Ile165. The enhanced rotation frequency of the side chain of Ile165 serves as a temperature-dependent revolving door. The increased frequency of door openings provides an opportunity for FINDY (**1**) to enter the pocket. The tight binding between DYRK1A and the selective inhibitors after transient heating suggests that the subsequent cooling to the physiological temperature stiffened the door movement, effectively packing the bulky molecules inside the pocket.

The slight difference in side chain orientation enables slow- and tight-binding inhibition because rotation of the side chains contribute to the stable protein–protein and protein–inhibitor interactions.^40–42^ For example, in cyclooxygenase-1 (COX-1), the reorientation of Leu531 allows for slow and tight binding with COX-1 inhibitors.^43^ For DYRK1A and FINDY (**1**), the reoriented conformations that are open to allow FINDY (**1**) to enter the binding site exist in a low-populated state at equilibrium. The lower population of FINDY (**1**)-binding-ready conformations could indicate a larger Δ*G*_BC(FINDY)_. The high energy barrier associated with Ile165 side-chain rotation may contribute to the small *k*_off_ value and tight binding of FINDY (**1**).

This study provides insight into the reasons why FINDY (**1**) inhibits DYRK1A but not DYRK1B in cultured cells and the *in vivo Xenopus* model, which remained elusive in our previous study.^28^ The dissociation rate constant of FINDY (**1**) for DYRK1A was approximately 22-fold smaller than that for DYRK1B. The slower dissociation rate results in longer residence time of the FINDY (**1**)–target complex. Consequently, when the effective concentration of FINDY (**1**) was reduced by its aggregation, adsorption, and metabolism in the cellular systems, the inhibitory potency was sustained for DYRK1A but not for DYRK1B. Thus, the dissociation rate may underlie the selectivity of FINDY (**1**) between DYRK1A and DYRK1B. The large difference between the dissociation rate constants is closely related to that between the association rate constants. The association rate constant of FINDY (**1**) for DYRK1A was smaller than that for DYRK1B, suggesting that DYRK1A requires greater free energy input than DYRK1B to reach the binding-competent state for FINDY (**1**), which is consistent with the melting temperature of DYRK1A being higher than that of DYRK1B.^44^ Although kinases possess a conserved ATP-binding pocket, the thermodynamics of the pocket is governed by dynamics-based allostery throughout the kinase domain,^45^ the thermal stability of which varies between kinases.^46^ Thus, the thermodynamics of the kinase domain affects the association and dissociation kinetics of inhibitors. The association rate is also the determinant related to target occupancy.^47^ Therefore, the balance between association and dissociation rates needs to be taken into account to develop a potent inhibitor.

*dp*-FINDY (**2**) exhibited remarkedly slow-binding kinetics for DYRK1A, in which its inhibitory potency did not reach a plateau even after 48 hr of pre-incubation. On the other hand, *dp*-FINDY (**2**) suppressed the intramolecular autophosphorylation selectively catalyzed by DYRK1A in the non-native state when DYRK1A was expressed for 30 min in the presence of *dp*-FINDY (**2**) in the reconstituted cell-free system.^33^ A large gap exists between these two experiments with regard to the time required for *dp*-FINDY (**2**) to exert its inhibitory effect. In the latter cell-free expression system, a ribosome-bound cotranslational folding intermediate could exist in the non-native state for a relatively extended period of time dependent on the translation rate.^48^ In addition, it has been reported that ribosomes trap and sustain a partially folded intermediate on the surface during translation.^49^ The cotranslational folding intermediate of DYRK1A, which is possibly homologous to the heat-evoked conformation, could bind rapidly to *dp*-FINDY (**2**). Upon translation termination, the complex with *dp*-FINDY (**2**) is released from the ribosome and converged to the stable bound state. Thus, in accordance with the conformational states of a target protein, such as cotranslational folding and chaperone-assisted refolding processes, a non-native-state-targeted inhibitor may bind rapidly.

Given the structural similarity between kinases, the slow association/dissociation-related selectivity improvement observed with the combination of DYRK1A/1B and FINDY (**1**) could be applied to other kinases. The structure–activity relationship study of FINDY (**1**) suggests that a slight alteration in chemical structure impacts inhibition kinetics. The difference of only one methyl group between the selective inhibitor (RD0448 (**11**)) and the canonical inhibitor (RD0562 (**12**)) leads us to believe that chemical libraries of kinase inhibitors and their structural derivatives, which have been synthesized with great effort in the long history of medicinal chemistry, contain non-native-state-targeted inhibitors. Slow- and tight-binding inhibitors have been missed in the conventional screening assays using *K*_D_ and IC_50_ values as indicators.^50^ In addition, there is concern that an inhibitor that appears highly selective may exhibit slow- and tight-binding to off-target kinases. The temperature vaulting method could address these concerns. This method may be applicable to other proteins with conformation, while further research is necessary. The advancement in accurate predictions of protein thermal fluctuation can facilitate the identification of a functional conformation in the low-populated non-native state as a drug target.

## EXPERIMENTAL SECTION

### Chemical compounds

Synthesis of FINDY (**1**), RD0447 (**3**), RD0392 (**5**), RD0448 (**11**), and RD0561 (**12**) was previously described.^28^ Synthesis of *dp*-FINDY (**2**) and its derivatives shown in Figures S4 and S10B was previously described.^33^ Synthesis of RD1408 (**4**), RD0562 (**13**), and RD1486 (**14**) is described in the Supporting Information. Harmine (**6**) (product no.: H0001) and staurosporine (**7**) (product no.: T4000) were obtained from Tokyo Chemical Industry Co., Ltd. (Tokyo, Japan). AZ191 (**8**) (cat. no.: HY-12277), GSK-626616 (**9**) (cat. no.: HY-105309), and leucettine L41 (**10**) (cat. no.: HY-117049) were obtained from MedChemExpress (New Jersey, USA).

### Temperature measurement

A portable 12-ch temperature measurement logger (BTM-4208SD) (Satotech, Sato Shouji Inc., Kanagawa, Japan) equipped with ultra-fine coated thermocouple T-type (TJT-CM32FP; Satotech) was used to measure solution temperatures (15 μL) in a PCR tube set on a thermal cycler (Biometra TAdvanced 96 SG, Analytik Jena GmbH, Germany).

### Preparation of recombinant proteins

The expression vector of His-ZZ-DYRK1A kinase domain, pCold-I-Km-ZZ-DYRK1A, was previously described.^35^ A DNA fragment coding Avi-tag peptide (GLNDIFEAQKIEWHE) was inserted at the carboxy terminus of DYRK1A kinase domain in pCold-I-Km-ZZ-DYRK1A to produce pCold-I-Km-ZZ-DYRK1A-Avitag. A DNA fragment of the kinase domain of DYRK1B (78–442; GenBank accession NM_004714) with stop codon was inserted into the *Bam*HI/*Afl*II site on pCold-I-Km-ZZ,^35^ producing pCold-I-Km-ZZ-DYRK1B. All constructs were verified by DNA sequencing (Eurofins Genomics K.K., Tokyo, Japan). The recombinant proteins were prepared as described previously.^35^ In brief, Rosetta 2 (DE3) pLysS competent cells (Novagen, Merck KGaA, Darmstadt, Germany) were transformed with the expression vectors. The cells were incubated with 0.5 mM of IPTG (Nacalai Tesque, Kyoto, Japan) at 6 ℃ for 16 hr, then suspended in lysis buffer (50 mM Tris-HCl, pH 7.5, 500 mM NaCl, 10% glycerol, 20 mM imidazole) and sonicated in the presence of lysozyme from egg white (Nacalai Tesque) and DNase I from bovine pancreas (Sigma-Aldrich, Merck). The clarified supernatants were loaded onto a AKTA start chromatography system equipped with a HiTrap TALON Crude column (Cytiva, Massachusetts, USA). The bound protein was eluted with imidazole (Nacalai Tesque). The His-ZZ-DYRK1A-Avitag protein was biotinylated using a BirA biotin–protein ligase standard reaction kit (Avidity LLC, Colorado, USA) according to the manufacturer’s instructions.

### In vitro kinase assay

For the temperature vaulting method, recombinant His-ZZ-DYRK1A kinase domain (350 ng) or His-ZZ-DYRK1B kinase domain (350 ng) was mixed in a total volume of 15 μL of kinase buffer (5 mM MOPS, pH 7.2, 2.5 mM β-glycerol-phosphate, 5 mM MgCl_2_, 1 mM EGTA, 0.4 mM EDTA, 0.25 mM DTT) containing the compounds at the concentrations indicated on the horizontal axes of Figures 2B–O, 5B–I, S4, and Figure S10B,C. The mixture in a well of a 96-well plate was transiently heated in the thermal cycler. Thereafter, 10 μL of a mixture containing BSA (Sigma-Aldrich, Merck), ATP (Promega Corporation, Wisconsin, USA), and DYRKtide peptide (RRRFRPASPLRGPPK) in kinase buffer was added into each well, resulting in final concentrations of 0.1 mg·mL^−1^, 20 μM, and 20 μM, respectively. Unless otherwise noted, the kinase reaction was performed at 37 ℃ for 40 to 90 min. The consumption of ATP was measured using a Kinase-Glo Luminescent Kinase Assay or ADP-Glo Kinase Assay (Promega Corporation). Luminescence was measured using a TriStar^2^S LB942 microplate luminometer (Berthold Technologies, Bad Wildbad, Germany). In the conventional kinase assay, the concentrations indicated on the horizontal axes of Figures 1C, 2B–O, 4A–G, 5B–N, S4, and Figure S10B,C are the final concentrations in the reaction mixture with BSA/ATP/DYRKtide. Thus, the final compound concentrations in the reaction mixture with BSA/ATP/DYRKtide for the temperature vaulting method are three-fifths of the final compound concentrations in the reaction mixture for the conventional assay. In the pre-incubation experiments except for Figure 5J,M,N and Figure S7C,D, the recombinant kinases and the compounds at the concentrations indicated on the horizontal axes of Figures 4A–G, 5K–L and Figure S7A,B were mixed with kinase buffer in PROKEEP protein low binding tubes (Watson Co., Ltd., Tokyo, Japan), which were then incubated at 37 ℃ for the indicated periods. In the pre-incubation experiments shown in Figure 5J,M,N and Figure S7C,D, a 96-well PCR plate was used to incubate the mixtures because of the short time interval. To 15 μL of the pre-incubated mixtures, 10 μL of the BSA/ATP/DYRKtide mixture was added prior to the kinase reaction for 15 min.

### Determination of Z′-factor

To assess the effect of transient heating on the quality of the assay, Z′-factor scores were determined as described previously.^51^ The Z′-factor scores were calculated with the actual luminescence values (raw data) from the kinase assay shown in Figure S3.

### Removal of unbound inhibitors from the protein mixture

His-ZZ-DYRK1A (final conc.: 23 μg·mL^−1^) or His-ZZ-DYRK1B (final conc.: 46 μg·mL^−1^) proteins and the inhibitors were mixed in kinase buffer (540 μL). The mixture was dispensed into a 96-well plate at 15 μL per well and heated at the indicated temperature for 20 sec. The mixtures were collected and subjected to a PD-10 desalting column (Cytiva) equilibrated with kinase buffer containing the solvent DMSO. The eluates were subjected to the kinase reaction.

### Thermal shift assay

The thermal stability of the kinase domain of DYRK1A in the presence or absence of the inhibitors was measured as previously described.^52^ In brief, 10 μM recombinant DYRK1A kinase domain (residues 127–485),^33^ 25 mM HEPES (pH 7.5), 500 mM NaCl, and 0.5 mM TCEP containing 100 μM of the inhibitor were reacted with SYPRO orange Protein Thermal Shift Dye (ThermoFisher Scientific, Inc., Massachusetts, USA) in 20 μL of reaction volume. Fluorescence intensities were monitored every 4 sec with a ramp rate of 0.02 ℃·sec^−1^ from 25 ℃ to 95 ℃ by using a QuantStudio 6 PCR system (ThermoFisher Scientific, Inc.). The fluorescence signal was fitted to a first-derivative curve to identify the melting temperatures. Three independent data sets were obtained.

### Surface plasmon resonance (SPR)

A Biacore X100 plus package (Cytiva) was used to monitor binding kinetics. All immobilization and binding experiments were performed at 25 ℃ in 10 mM HEPES, pH 7.4, 150 mM NaCl, 3 mM EDTA, 0.05% Tween-20, and 5% DMSO (v/v). A solution of the biotinylated His-ZZ-DYRK1A-Avitag protein (0.7 mg·mL^−1^) was injected on a Sensor Chip SA (Cytiva). The protein was immobilized at a density between 8,000 and 10,000 RU. Prior to analysis, solvent (DMSO) calibration and double referencing subtractions were done. “Single Cycle Kinetic” analysis was used to generate binding sensorgrams. Affinity and binding kinetics parameters were determined by global fitting to a two-state binding model within the Biacore X100 Evaluation Software ver. 2.0.1 (Cytiva). The dissociation rate constant for the two-state model was defined as *k*_off_ = *k*_d1_*k*_d2_/(*k*_a2_ + *k*_d2_).

### Molecular dynamics simulation

The kinase domain (residues 133–481) structure of DYRK1A was taken from the native (PDB ID: 7FHT). The missing residues were modeled by MODELLER.^53^ The simulation system consisted of the kinase-domain structure and solvent molecules of 100 mM KCl solution, which was confined in a truncated octahedron box with a periodic boundary condition. The force field of ff14SB was applied for the protein. TIP3P water model was used. We performed energy minimization, and MD simulation for the simulation system to heat from 1 K to 298.15 K for 1 nsec and equilibrate under 1 bar for 1 nsec. Starting from the equilibrated system, production runs of MD simulations were conducted at the targeted temperatures of 25, 40, 55, 70, 85 and 100 ℃. In each run, the 2-μsec trajectory was simulated four times (total of 8 μsec at the one temperature). The snapshots were saved every 100 psec during the production runs to be provided for RMSD analysis and PCA. The software AmberTools20 and Amber20 were used.^54^ The RMSD values were calculated for the main-chain atoms except for the flexible N-terminal region (residues 133–143) of snapshots after superimposition onto the initial native structure. The main-chain dihedral angles of the snapshots were inputted as feature vectors for PCA. The plane constructed by PC1 and PC2 axes was used to show the conformational distribution of snapshots.

### Analysis of the dose–response curves

The dose–response curves in Figures 2B–O, 4A–G, 5B–N, S4, and Figure S10B,C were fitted to a four-parameter logistic curve (variable slope) for Hill slope determination, from which the IC_50_ values were calculated using GraphPad Prism ver. 8.0 (GraphPad Software, California, USA). The IC_50_ values are summarized in Tables S1– S4.

### Analysis of the pre-incubation titration curves

The pre-incubation titration curves in Figure S7 were analyzed according to a previously reported method.^36,37^ Binding kinetics parameters were determined by global fitting to eq 15 using GraphPad Prism ver. 8.0.

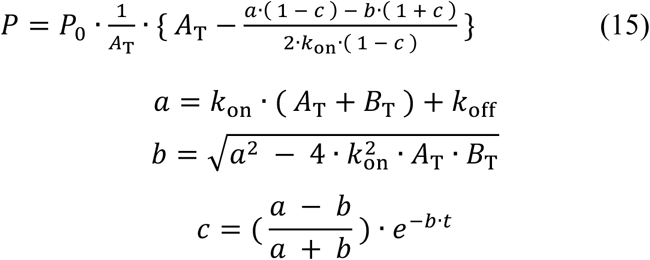

where *k*_on_ and *k*_off_ are the association and dissociation rate constants, respectively. *A*_T_ and *B*_T_ are the total concentrations of the kinase protein and the inhibitor, respectively. *t* is the pre-incubation time. *P* is the relative enzymatic activity, and *P*_0_ is the relative enzymatic activity when *B*_T_ = 0. The fitting curves are shown in Figure S7. The kinetics parameters are summarized in Table 1.

## SUPPORTING INFORMATION

The supporting information describes synthesis of compounds, temperature measurement in the thermal cycler, Z′-factor determination, additional kinase assays, PCA of MD simulation, surface plasmon resonance analysis, thermal shift assay, and summary of IC_50_ values (PDF).

## NOTES

The authors declare the following competing financial interest: I.K., G.F., and K.U. are co-inventors on a patent owned by Shinshu University, which covers the use of temperature vaulting methods for *in vitro* assays: WO/2022/118941 and PCT/JP2021/044375. The co-inventors license for-profit companies under non-disclosure and non-exclusive license agreements. The other authors declare that they have no competing interests.

## ACKNOWLEDGMENTS

We thank all the laboratory members for their critical comments and discussion. We thank Ms. Oda and members of the Instrumental Center of Shinshu University for assistance with the surface plasmon resonance measurements. We thank Uni-edit (https://uni-edit.net/) for editing the manuscript. This work was supported by JSPS KAKENHI grant numbers JP24K01962 (I.K.), JP19H02856 (I.K.), JP18H04568 (I.K.), JP17H05678 (I.K.), JP16H05926 (I.K.), and JP24K02184 (T.U.), JP20H03388 (T.U.); AMED grand numbers JP23ym01268113 (I.K.), JP24ym0126813 (I.K.), and JP23nk0101646 (I.K.); the Cooperative Research Project of Research Center for Biomedical Engineering (I.K., T.H.); TAKEUCHI Scholarship Foundation (I.K.); Kobayashi Foundation (I.K.); the Life Science Foundation of JAPAN (I.K.); Tokyo Biochemical Research Foundation (I.K.); and Mochida Memorial Foundation for Medical and Pharmaceutical Research (I.K.).

## CONTRIBUTIONS

I.K., S.S., and K.U. contributed to conceptualization of the project. I.K. and G.F. obtained initial proof-of-concept data. I.K. and S.S. designed experiments. S.S., N.F., N.K., and G.F. performed kinase assays. K.U. conducted MD simulation. S.S. performed theoretical calculations and modeling. M.K. and T.U. performed thermal shift assays. M.Y. performed surface plasmon resonance measurements. N.K., N.F., S.S., and M.Y. prepared recombinant proteins. D.N., T.N., and T.H. synthesized small molecules. I.K., S.S., and K.U. prepared the manuscript with input from all authors.

