## Supporting Information for "Temperature vaulting: A method for screening of slow- and tight-binding inhibitors that selectively target kinases in their non-native state"

### Table of Contents


**Synthesis of chemical compounds** S3–S15

General Remarks

a) Synthesis of RD1408 (**4**)

b) Synthesis of RD0562 (**12**)

c) Synthesis of RD1486 (**13**)

NMR spectra of compounds

**Tables** S16–S19

Table S1. Summary of $IC_{50}$ values for the indicated inhibitors calculated from the dose–response curves in Figures 2B–E,G–O (DYRK1A), 5B–N and S10C (DYRK1B)

Table S2. Summary of $IC_{50}$ values for the indicated inhibitors calculated from the pre-incubation titration curves (DYRK1A) in Figure 4A–C and 4F,G

Table S3. Summary of $IC_{50}$ values for the indicated inhibitors calculated from the pre-incubation titration curves (DYRK1B) in Figure 5J–N

Table S4. Summary of $IC_{50}$ values for the indicated inhibitors calculated from the dose–response curves in Figures S4 (DYRK1A) and S10B (DYRK1B)

**Figures** S20–S30

Figure S1. Actual temperature change of a solution in the thermal cycler

Figure S2. Validation of the stability and reproducibility of the temperature change in the 96 wells of the thermal cycler

Figure S3. Robustness of the kinase assay with the temperature vaulting method

Figure S4. Inhibitory potency of the FINDY derivatives in the temperature vaulting method

Figure S5. FINDY (**1**) and *dp*-FINDY (**2**) did not inhibit the DYRK1A protein that was once denatured and then refolded

Figure S6. Analysis of the DYRK1A kinase domain complexed with the inhibitors

Figure S7. Inhibition of the DYRK1A/1B proteins by either FINDY (**1**) or *dp*-FINDY (**2**) under pre-equilibrium conditions

Figure S8. The association/dissociation kinetics of RD0392 (**5**) for the DYRK1A protein

Figure S9. Thermal stability of the DYRK1A kinase domain in the presence of the inhibitors

Figure S10. The transient-heating-dependent inhibition of the DYRK1B protein

**Supplementary references** S31

**Synthesis of chemical compounds**

**General Remarks**

Unless otherwise noted, all reactions were carried out under an argon atmosphere. Reactions were monitored by thin-layer chromatography (TLC) using E. Merck Kieselgel 60 N $F_{254}$ (0.25 mm thickness) and visualization was conducted using UV light (254 nm), *p*-anisaldehyde, and dinitrophenol stains. Column chromatography was conducted on a Yamazen automated flash chromatography system with a Universal Premium column ($SiO_2$, 30 μm sphere, 16 g [M size] or 40 g [L size]). Gel permeation chromatography was conducted on a YMC LC-Forte/R preparative HPLC system with YMC GPC T-2000 column (10 μm sphere, 22.1 mmΦ × 60 mm). Silica gel filtration was conducted on a Biotage Isolute® Si II prefilled column (200 mg/3 mL or 2 g/6 mL). A Prefilled Celite® column was purchased from Biotage AB (Isolute® HM-N Celite® pad). Membrane filtration was conducted on Cytiva Whatman Puradisc 12 (PVDF, pore size: 0.2 μm, 13 mmΦ). Melting points (mp) were measured with an OptiMelt® Automated melting point apparatus (Stanford Research Systems, Inc.) and were uncorrected. $^{1}$H and $^{13}$C NMR spectra were obtained from measurements at ambient temperature on a JEOL JNM-ECS400 (395.9 MHz for $^{1}$H NMR; 99.5 MHz for $^{13}$C NMR) spectrometer. Chloroform-*d*$_1$ ($CDCl_3$) containing 0.05% tetramethylsilane (TMS) (99.8% D, Euriso-Top SAS) and dimethylsulfoxide-*d*$_6$ (DMSO-*d*$_6$, 99.8% D, Euriso-Top SAS) were used as a solvent for NMR measurements at ambient temperature. Chemical shifts (δ) are given in parts per million (ppm) downfield from TMS (δ 0.00 ppm for $^{1}$H NMR in $CDCl_3$) or the solvent peak (δ 77.0 ppm for $^{13}$C NMR in $CDCl_3$, δ 2.50 ppm for $^{1}$H NMR in DMSO-*d*$_6$, and δ 39.52 ppm for $^{13}$C NMR in DMSO-*d*$_6$) as an internal reference with coupling constants (*J*) in hertz (Hz). The abbreviations s, d, t, q, br, and m signify singlet, doublet, triplet, quartet, broad, and multiplet, respectively. Infrared (IR) absorption spectra were measured by an attenuated total reflection (ATR) method on a Shimadzu IR Prestige-21 spectrometer equipped with a PIKE Technologies MIRacle™ single reflection ATR optical attachment. The absorption bands are given in wavenumber ($cm^{-1}$). High-resolution mass spectra (HRMS) were measured on a Thermo Fisher Scientific Exactive Plus™ ESI-Orbitrap™ mass spectrometer or a Thermo Fisher Scientific Q Exactive™ ESI-Orbitrap™ mass spectrometer under the positive electrospray ionization ($ESI^+$) or a Bruker micrOTOF mass spectrometer under negative electrospray ionization ($ESI^-$) conditions.

4-Hydroxy-3-iodobenzaldehyde was purchased from Angene International Limited. 4-Methoxy-3-iodobenzaldehyde was purchased from BLD Pharmatech Ltd. (Methyldiphenylsilyl)acetylene, bis(triphenylphosphine)palladium(II) dichloride

($PdCl_2(PPh_3)_2$), rhodanine, and thiohydantoin were purchased from Sigma–Aldrich Japan. Potassium fluoride (KF) was purchased from Kanto Chemical Co. Inc. Ammonium acetate ($NH_4OAc$) was purchased from FUJIFILM Wako Pure Chemical Corporation. Trimethylsilylacetylene, 3,4-methylenedioxybenzaldehyde, 3-hydroxy-4-methoxybenzaldehyde, 3-ethoxy-4-methoxybenzaldehyde, and 3,4-dimethoxybenzaldehyde were purchased from Tokyo Chemical Industry Co., Ltd. 4-Methoxybenzaldehyde, copper iodide (CuI), tetrahydrofuran (THF), triethylamine ($Et_3N$), diisopropylamine ($Et_2NH$), ethyl acetate (AcOEt), *n*-hexane, diethyl ether ($Et_2O$), chloroform ($CHCl_3$), dichloromethane ($CH_2Cl_2$), methanol (MeOH), acetonitrile (MeCN), acetic acid (AcOH), ammonium chloride ($NH_4Cl$), and anhydrous sodium sulfate ($Na_2SO_4$) were purchased from Nacalai Tesque, Inc. These reagents were used without further purification. 4-Methoxy-3-((methyldiphenylsilyl)ethynyl)benzaldehyde (**S1**) was prepared according to the reported procedure.[1]

**Scheme S1. Synthetic schemes of compounds.**

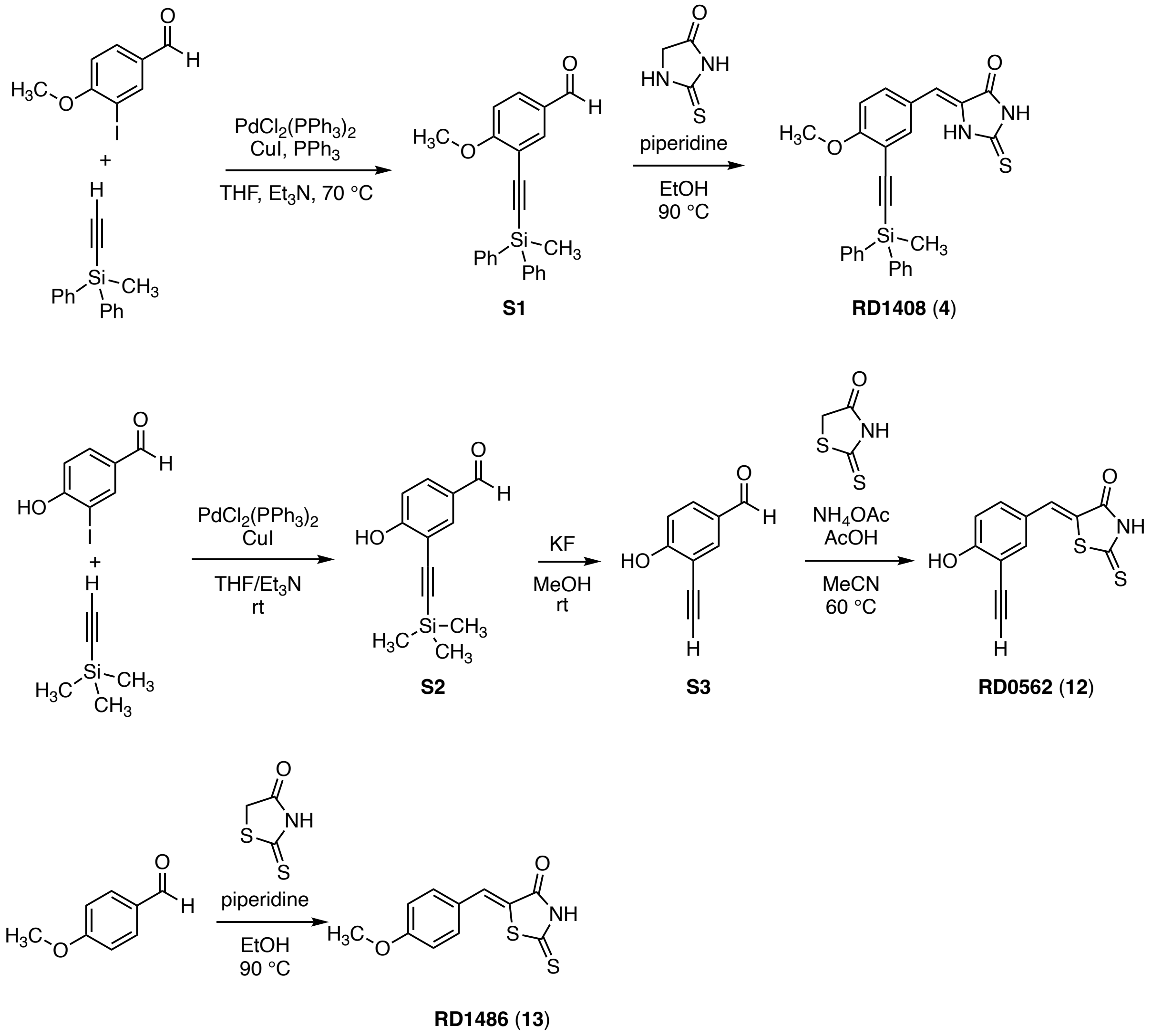

**a) Synthesis of RD1408 (4)**

*(Z)-5-(4-Methoxy-3-((methyldiphenylsilyl)ethynyl)benzylidene)-2-thioxoimidazolidin-4-one (4;* ***RD1408****)*

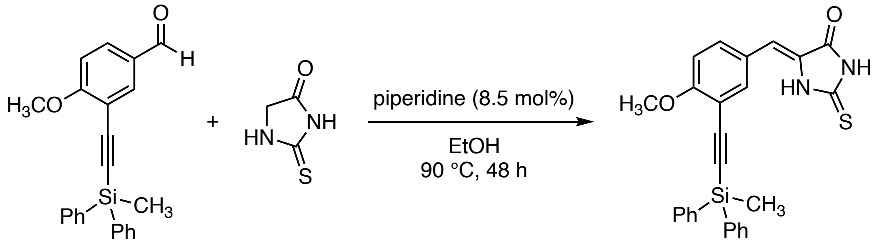


**S1** (1.1 equiv)  **RD1408** (**4**), 47%

In a 4 mL capped vial with a magnetic stirring bar were sequentially added thiohydantoin (232 mg, 2.00 mmol, 1 equiv), 4-methoxy-3-((methyldiphenylsilyl)ethynyl)benzaldehyde (**S1**, 756 mg, 2.12 mmol, 1.1 equiv), EtOH (3 mL), and piperidine (17 µL, 0.17 mmol, 8.5 mol%). The mixture was stirred for 48 h at 90 °C and then cooled to room temperature. After concentration, to the residue were added $Et_2O$ (1 mL) and brine (1 mL). The mixture was extracted with AcOEt (5 mL × 3). The combined organic extract was dried over $Na_2SO_4$. After filtration, the filtrate was concentrated under reduced pressure. The residue was diluted with $Et_2O$ and filtered through a silica-gel (2 g cartridge). After removal of the solvent in vacuo, to the residue were added $Et_2O$ (1 mL) and *n*-hexane (1 mL). The resulting precipitate was filtered to afford (*Z*)-5-(4-methoxy-3-((methyldiphenylsilyl)ethynyl)benzylidene)-2-thioxoimidazolidin-4-one (**4**; **RD1408**, 431 mg, 0.948 mmol, 47.4%) as a light yellow solid; mp 121 °C (decomp.); TLC: $R_f$ = 0.40 ($CHCl_3$/MeOH = 20:1); $^1$H NMR (400MHz, $CDCl_3$) δ 9.56 (brs, 1H), 9.40 (brs, 1H), 7.75–7.69 (m, 4H), 7.60 (d, *J* = 2.4 Hz, 1H), 7.43–7.34 (m, 7H), 6.89 (d, *J* = 8.8 Hz, 1H), 6.62 (s, 1H), 3.92 (s, 3H), 0.77 (s, 3H); $^{13}C\{^1H\}$ NMR (100 MHz, $CDCl_3$) δ 176.5 (1C), 164.7 (1C), 161.8 (1C), 135.0 (2C), 134.6 (4C), 134.5 (1C), 132.2 (1C), 129.7 (2C), 127.9 (4C), 126.3 (1C), 124.7 (1C), 113.7 (1C), 113.5 (1C), 111.4 (1C), 103.0 (1C), 96.7 (1C), 56.1 (1C), –2.1 (1C); IR ($cm^{-1}$) 1732, 1651, 1593, 1487, 1369, 1279, 953, 804, 723, 696, 619; HRMS ($ESI^+$) *m/z* 477.1062 (477.1063 calcd for $C_{26}H_{22}N_2NaO_2{}^{32}SSi^+$, $[M+Na]^+$).

**b) Synthesis of RD0562 (12)**

*4-Hydroxy-3-((trimethylsilyl)ethynyl)benzaldehyde (**S2**)*

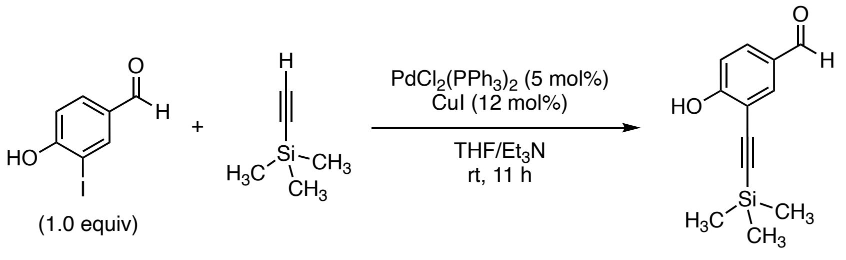


**S2** was prepared according to the reported procedure[2] with minor modifications.

To a 10 mL screw-capped vial with a magnetic stirring bar were sequentially added 4-hydroxy-3-iodobenzaldehyde (505 mg, 2.04 mmol, 1.02 equiv), $PdCl_2(PPh_3)_2$ (70.6 mg, 0.101 mmol, 5 mol%), CuI (48.2 mg, 0.253 mmol, 12 mol%), THF (10 mL), trimethylsilylacetylene (277 µL, 2.00 mmol), and $Et_3N$ (10 mL). The mixture was stirred for 11 h at room temperature and then the reaction mixture was quenched with saturated aqueous $NH_4Cl$ (5 mL). The mixture was extracted with AcOEt (5 mL × 3), and the combined organic extract was filtered through a silica-gel (2 g cartridge). The filtrate was concentrated under reduced pressure and the residue was purified by a silica-gel column chromatography (AcOEt/*n*-hexane) to afford 4-hydroxy-3-((trimethylsilyl)ethynyl) benzaldehyde (**S2**, 347 mg, 1.59 mmol, 79.4% yield) as a brown oil; TLC: $R_f$ = 0.48 (*n*-hexane/AcOEt = 7:1); $^1$H NMR (400 MHz, $CDCl_3$) δ 9.83 (s, 1H), 7.90 (d, $J$ = 2.4 Hz, 1H), 7.80 (dd, $J$ = 8.8, 2.4 Hz, 1H), 7.07 (d, $J$ =8.8 Hz, 1H), 0.29 (s, 9H) (the signal for the proton attached to the oxygen atom was not observed); $^{13}C\{^1H\}$ NMR (100 MHz, $CDCl_3$) δ 190.0 (1C), 161.7 (1C), 134.4 (1C), 132.0 (1C), 129.6 (1C), 115.3 (1C), 110.5 (1C), 102.1 (1C), 97.1 (1C), –0.2 (3C); IR (cm$^{-1}$) 1672, 1574, 1296, 1281, 1248, 1171, 1113, 1101, 839, 756; HRMS (ESI$^+$) *m/z* 241.0657 (241.0655 calcd for $C_{12}H_{14}NaO_2Si^+$, $[M+Na]^+$).

*3-Ethynyl-4-hydroxybenzaldehyde (**S3**)*

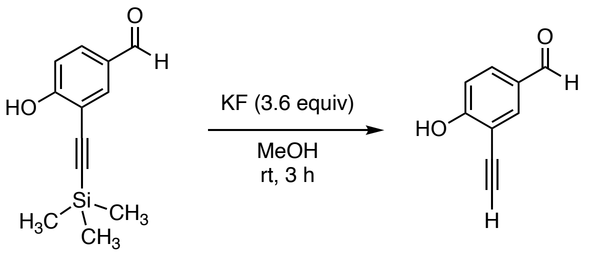


To a 20 mL round-bottom flask with a magnetic stirring bar were added KF (335 mg, 5.77 mmol, 3.6 equiv) and MeOH (5 mL), and then the mixture was stirred at room temperature until the KF dissolved in MeOH. To this was added a solution of 4-hydroxy-3-((trimethylsilyl)ethynyl)benzaldehyde (**S2**, 347 mg, 1.59 mmol) in MeOH (3 mL), and the reaction mixture was stirred for 3 h at room temperature. The reaction mixture was poured into brine (5 mL), and the mixture was extracted with $CH_2Cl_2$ (4 mL × 3). The combined organic extract was dried over $Na_2SO_4$. After filtration, the filtrate was concentrated under reduced pressure. The residue was dissolved in $CHCl_3$ (4 mL), and then insoluble solids were removed by filtration using a membrane filter. The filtrate was concentrated and purified by using a gel permeation chromatography ($CHCl_3$) to afford 3-ethynyl-4-hydroxybenzaldehyde (**S3**, 40.5 mg, 0.277 mmol, 17.4% yield) as a colorless solid; mp. 89.5 °C (decomp.); TLC: $R_f$ = 0.20 (*n*-hexane/AcOEt = 4:1); $^1$H NMR (400 MHz, $CDCl_3$) δ 9.85 (s, 1H), 7.94 (d, $J$ = 2.0 Hz, 1H), 7.83 (dd, $J$ = 8.8, 2.0 Hz, 1H), 7.09 (d, $J$ = 8.8 Hz, 1H), 6.48 (brs, 1H), 3.55 (s, 1H); $^{13}C\{^1H\}$ NMR (100 MHz, DMSO-$d_6$) δ 190.7 (1C), 162.1 (1C), 164.4 (1C), 136.3 (1C), 131.3 (1C), 128.3 (1C), 116.1 (1C), 110.0 (1C), 84.9 (1C); IR (cm$^{-1}$) 1661, 1591, 1566, 1503, 1287, 1258, 1179, 1111, 810, 704; HRMS (ESI$^-$) *m/z* 145.0297 (145.0295 calcd for $C_9H_5O_2^-$, $[M–H]^-$).

*(Z)-5-(3-Ethynyl-4-hydroxybenzylidene)-2-thioxothiazolidin-4-one (**12**; **RD0562**)*

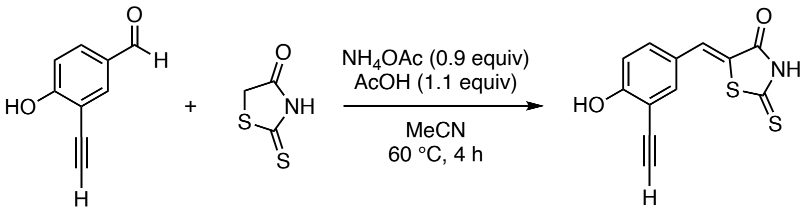


To a 4 mL screw-capped vial with a magnetic stirrer bar were sequentially added 3-ethynyl-4-hydroxybenzaldehyde (**S3**, 30.4 mg, 0.208 mmol, 1.1 equiv), $NH_4OAc$ (14.4 mg, 0.187 mmol, 0.9 equiv), rhodanine (26.2 mg, 0.197 mmol), MeCN (0.4 mL), and AcOH (12.3 µL, 0.215 mmol, 1.1 equiv). The reaction mixture was stirred for 4 h at 60 °C. After cooling to room temperature, to the reaction mixture was added $H_2O$ (1 mL). The resulting precipitate was collected by filtration and sequentially washed with $H_2O$ (2 mL) and *n*-hexane (2 mL) on the funnel. The solid was dried in vacuo to afforded (*Z*)-5-(3-ethynyl-4-hydroxybenzylidene)-2-thioxothiazolidin-4-one (**12**; **RD0562**, 45.4 mg, 0.174 mmol, 88.3% yield) as an orange solid; mp: 146.0 °C (decomp.); TLC: $R_f$ = 0.30 ($CHCl_3$/MeOH = 20:1); $^1$H NMR (400 MHz, DMSO-$d_6$) δ 11.0 (s, 1H), 7.60 (d, $J$ = 2.4 Hz, 1H), 7.55 (s, 1H), 7.46 (dd, $J$ = 9.2, 2.4 Hz, 1H), 7.05 (d, $J$ = 9.2 Hz, 1H), 4.30 (s, 1H) (the signal for the proton that is attached to the nitrogen atom was not observed); $^{13}$C{$^1$H} NMR (100 MHz, DMSO-$d_6$) δ 195.5 (1C), 169.7 (1C), 161.3 (1C), 136.7 (1C), 132.5 (1C), 131.3 (1C), 124.2 (1C), 122.4 (1C), 116.7 (1C), 110.5 (1C), 85.0 (1C), 79.5 (1C); IR (cm$^{-1}$) 1680, 1672, 1591, 1574, 1406, 1188, 1165, 1107, 914, 623; HRMS (ESI$^-$) *m/z* 259.9845 (259.9845 calcd for $C_{12}H_6NO_2{}^{32}S_2^-$, [M−H]$^-$).

**c) Synthesis of RD1486** (**13**)

*(Z)-5-(4-Methoxybenzylidene)-2-thioxothiazolidin-4-one (**13**; **RD1486**)*

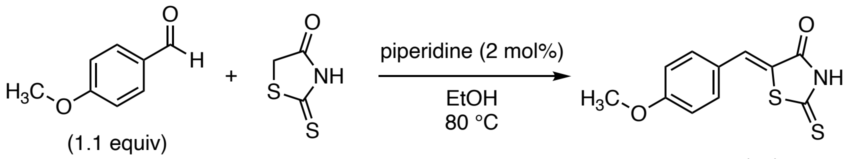


**RD1486** (**13**)

**RD1486** was prepared according to the reported procedure[3] with minor modifications.

To a 4 mL capped vial with a magnetic stirring bar were added rhodanine (240 mg, 1.80 mmol), EtOH (3 mL), 4-methoxybenzaldehyde (244 µL, 2.01 mmol), and piperidine (3 µL, 30 µmol, 2 mol%), and the reaction mixture was stirred for 16 h at 80 °C. After cooling to room temperature, the resulting precipitate was collected by filtration. The solid was sequentially washed with $H_2O$ (2 mL) and *n*-hexane (2 mL) on the funnel, and then dried in vacuo to afford (*Z*)-5-(4-methoxybenzylidene)-2-thioxothiazolidin-4-one (**13**; **RD1486**, 447 mg, 1.78 mmol, 98.7% yield) as a yellow solid; mp: 254 °C (decomp.); TLC: $R_f$ = 0.50 ($CHCl_3$/MeOH = 20:1); $^1$H NMR (400 MHz, DMSO-$d_6$) δ 7.62 (s, 1H), 7.57 (AA'BB', 2H), 7.11 (AA'BB', 2H), 3.83 (s, 3H) (the signal for the proton that is attached to the nitrogen atom was not observed); $^{13}$C{$^1$H} NMR (100 MHz, DMSO-$d_6$) δ 195.5 (1C), 169.5 (1C), 161.3 (1C), 132.7 (2C), 131.9 (1C), 125.5 (1C), 122.2 (1C), 115.1 (2C), 55.6 (1C); IR (cm$^{-1}$) 1682, 1582, 1557, 1508, 1258, 1236, 1013, 822, 679, 602; HRMS (ESI$^+$) *m*/*z* 273.9967 (273.9967 calcd for $C_{11}H_9NNaO_2{}^{32}S_2{}^+$, [M+Na]$^+$).

$^{1}H$ NMR (400 MHz) and $^{13}C\{^{1}H\}$ NMR (100 MHz) spectra of **RD1408** (**4**) ($CDCl_3$)

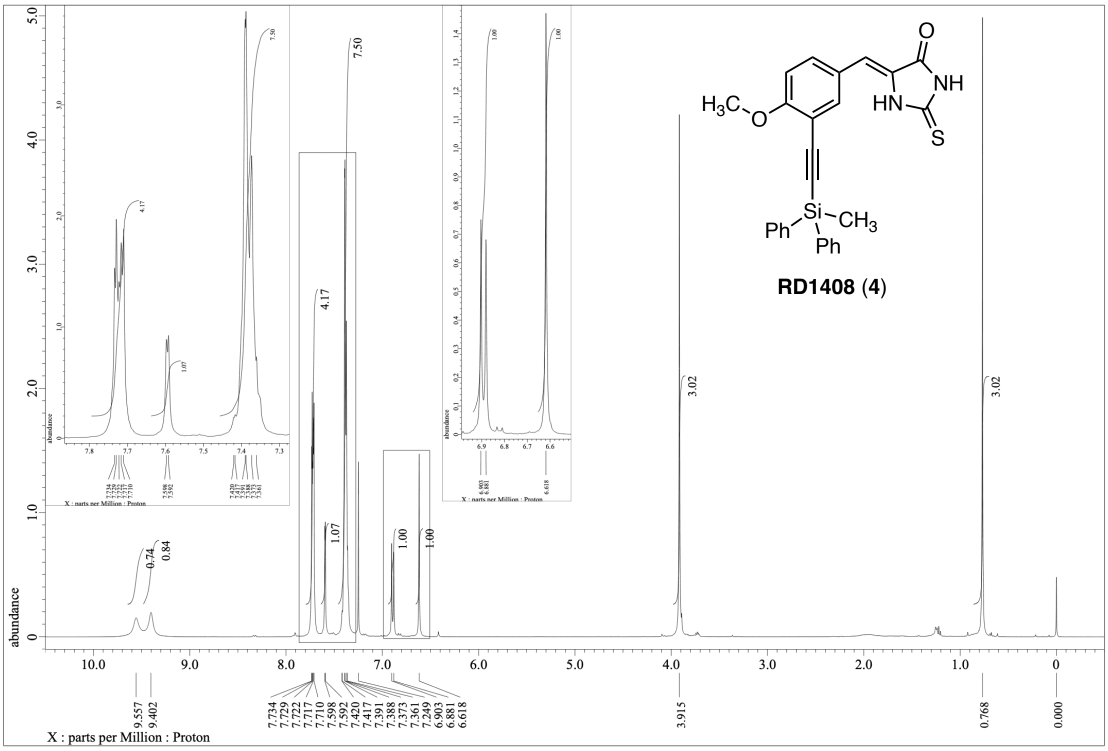


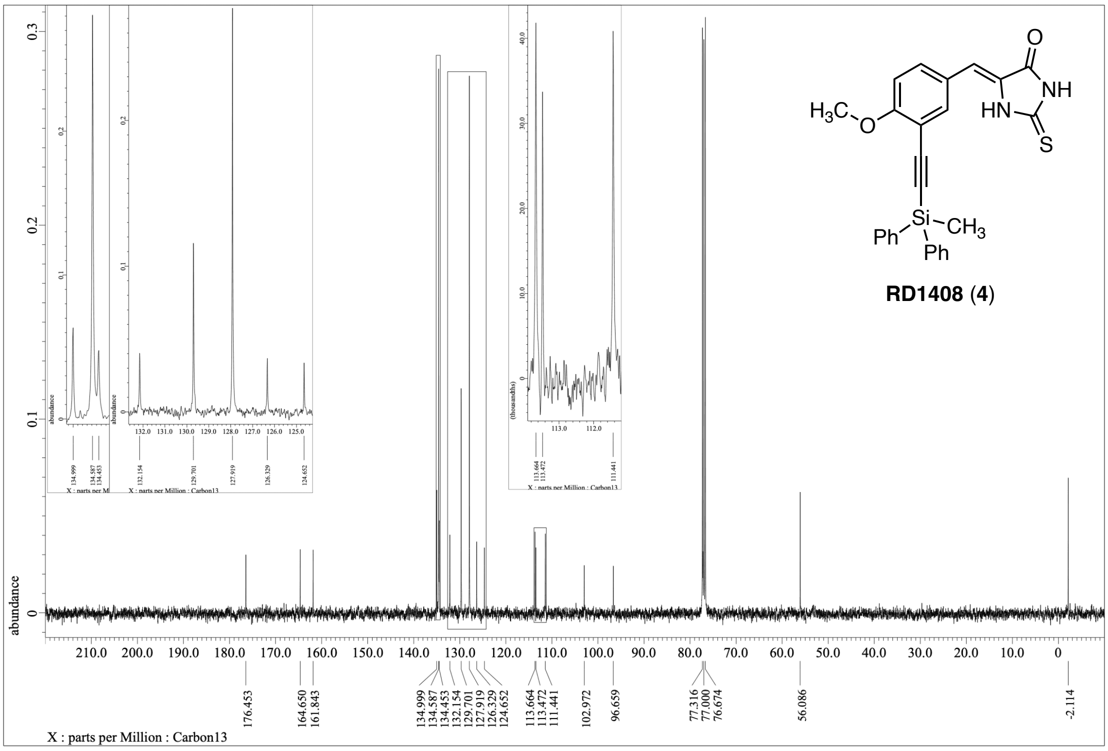

$^1H$ NMR (400 MHz) and $^{13}C\{^1H\}$ NMR (100 MHz) spectra of **S2** ($CDCl_3$)

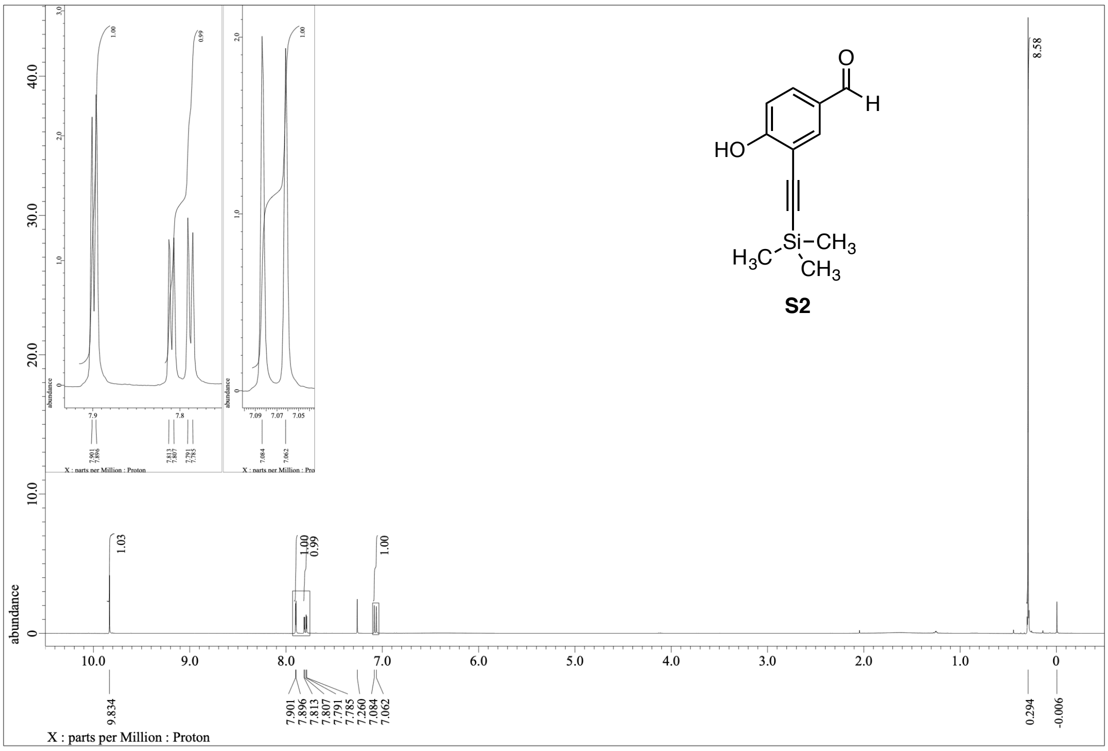


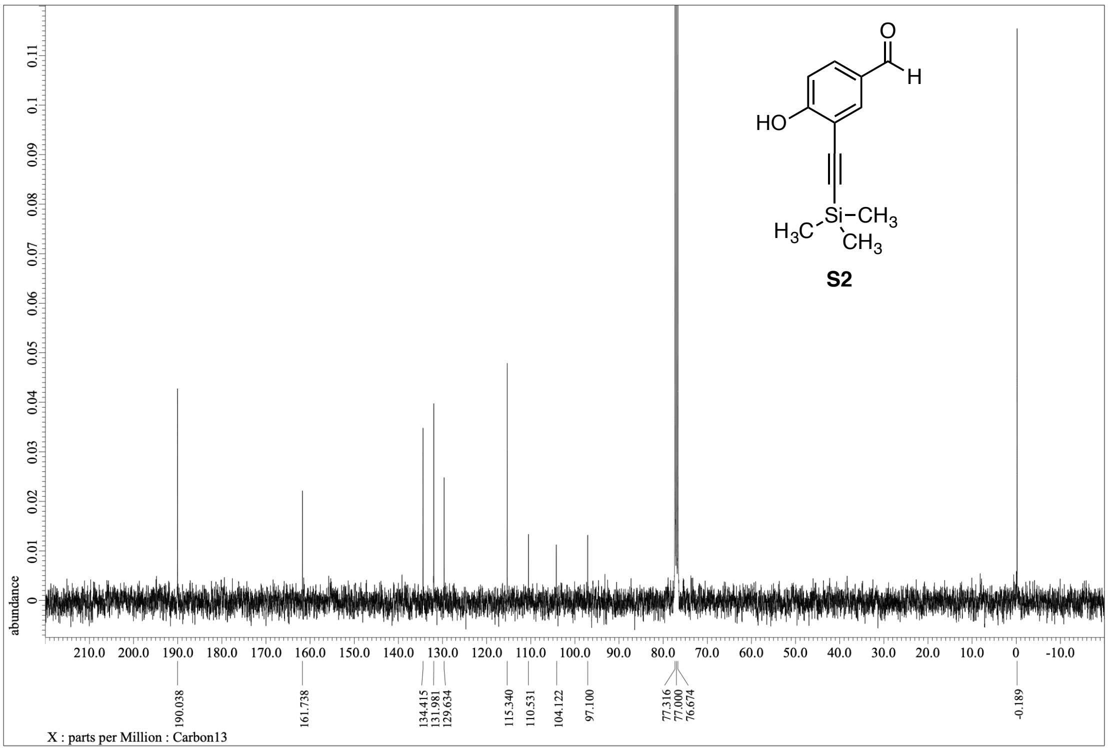

[1]H NMR (400 MHz, $CDCl_3$) and [13]C{[1]H} NMR (100 MHz, DMSO-$d_6$) spectra of **S3**

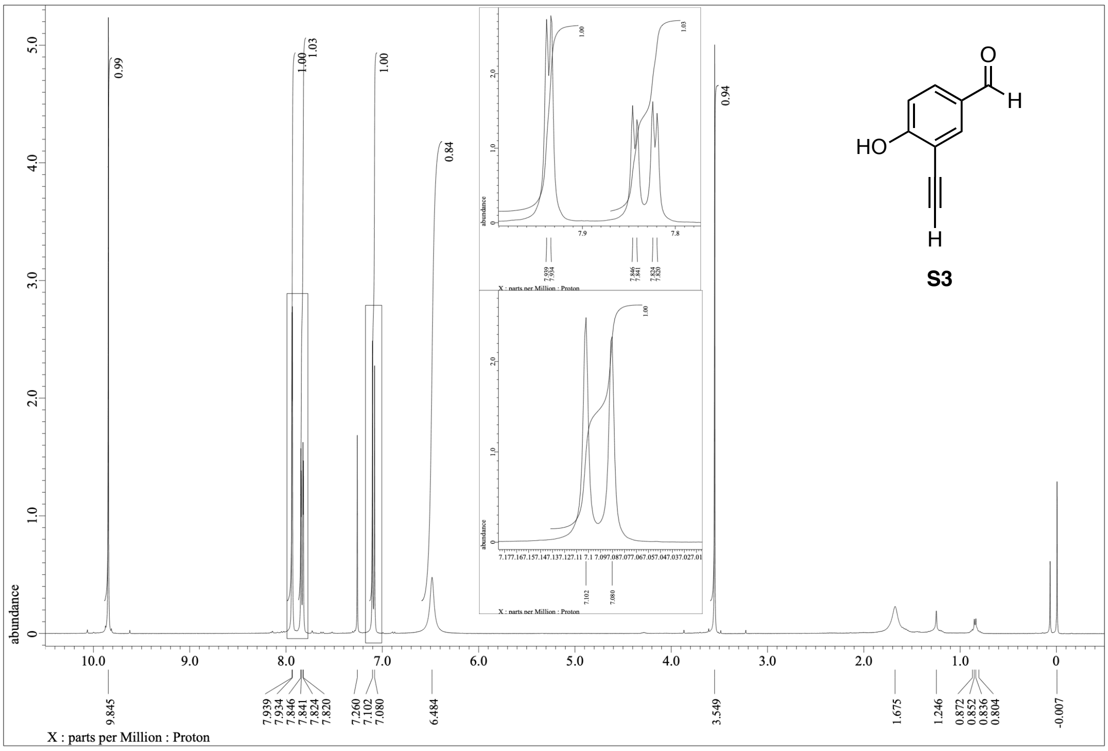


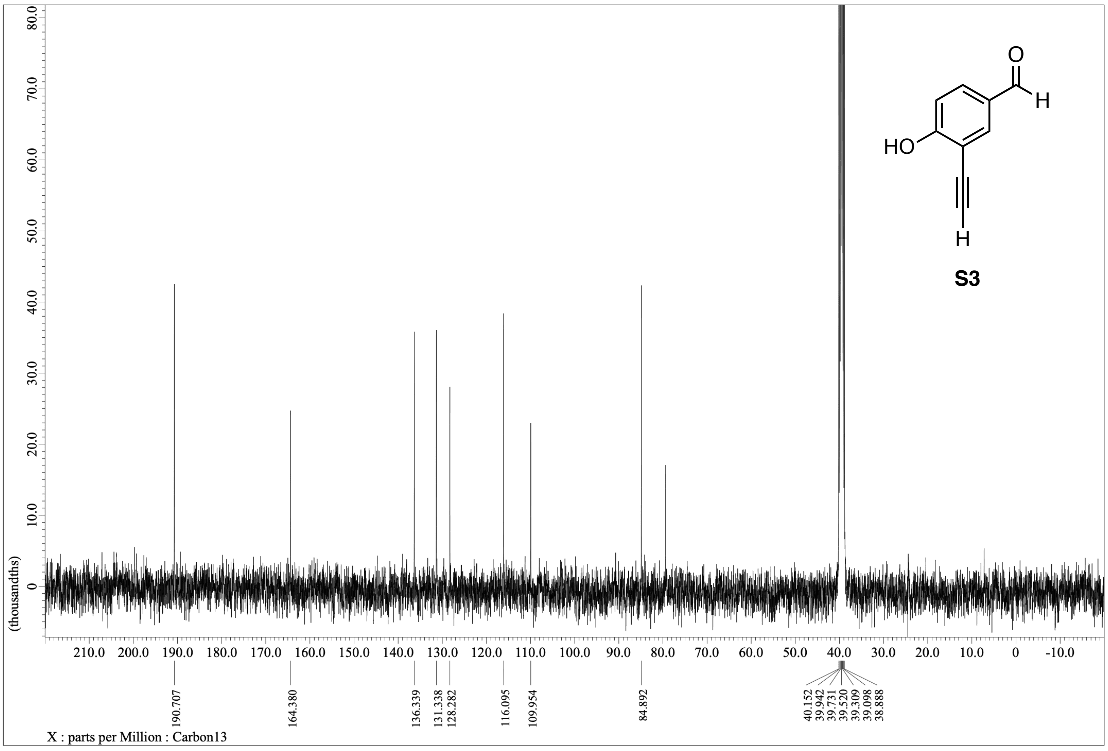

$^{1}$H NMR (400 MHz) and $^{13}$C{$^{1}$H} NMR spectra (100 MHz) of **RD0562** (**12**) (DMSO-$d_6$)

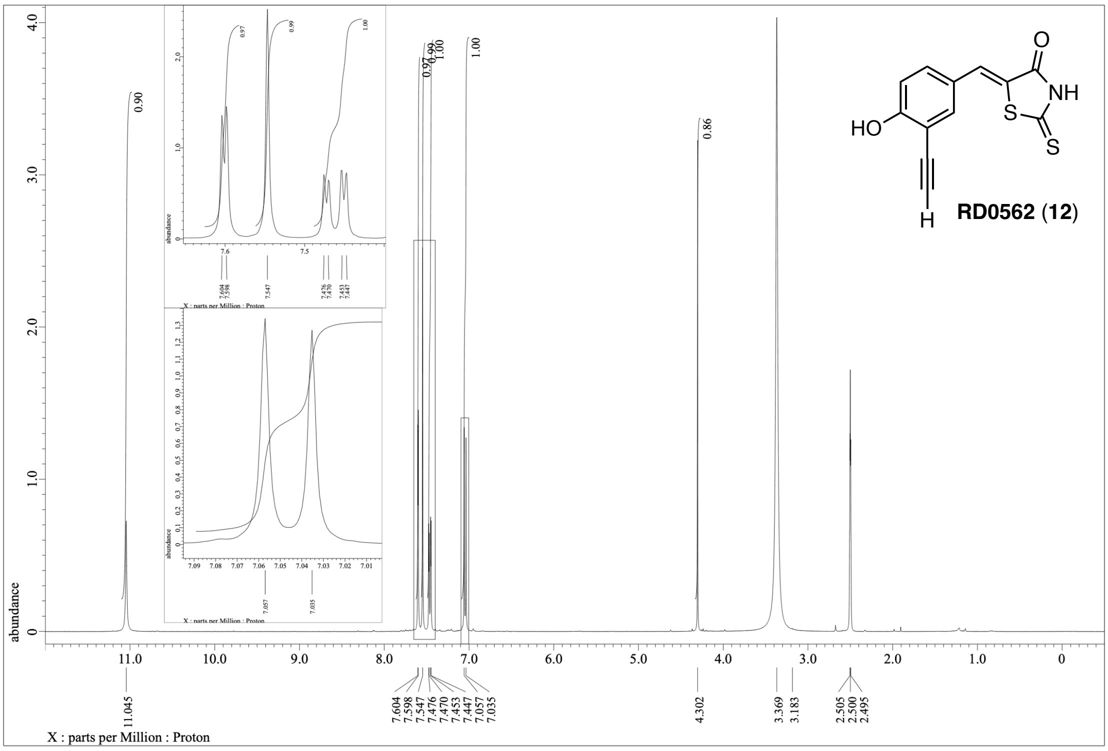


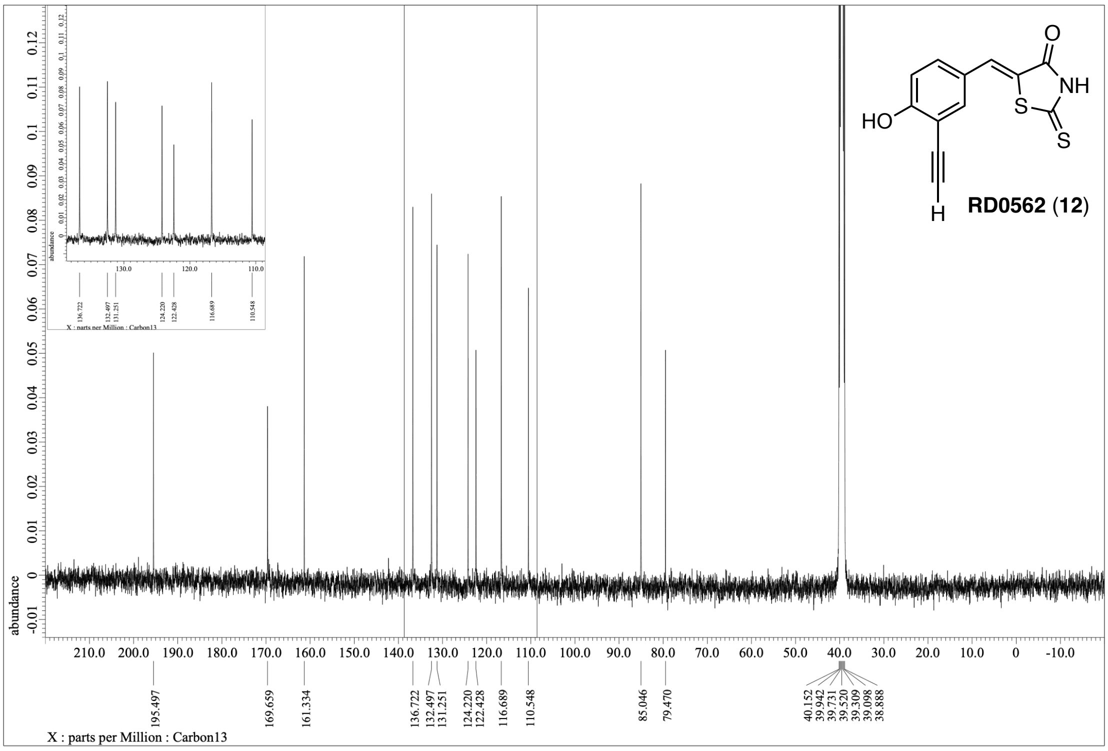

[1]H NMR (400 MHz) and [13]C{[1]H} NMR (100 MHz) spectra of **RD1486** (**13**) (DMSO-$d_6$)

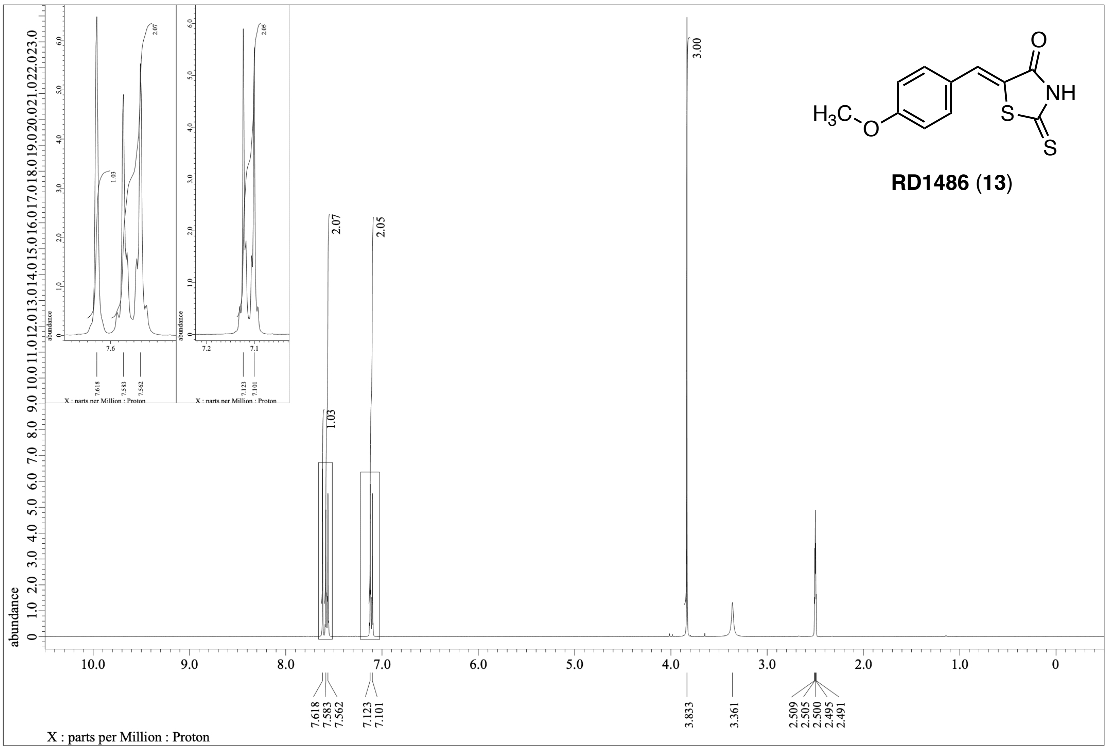


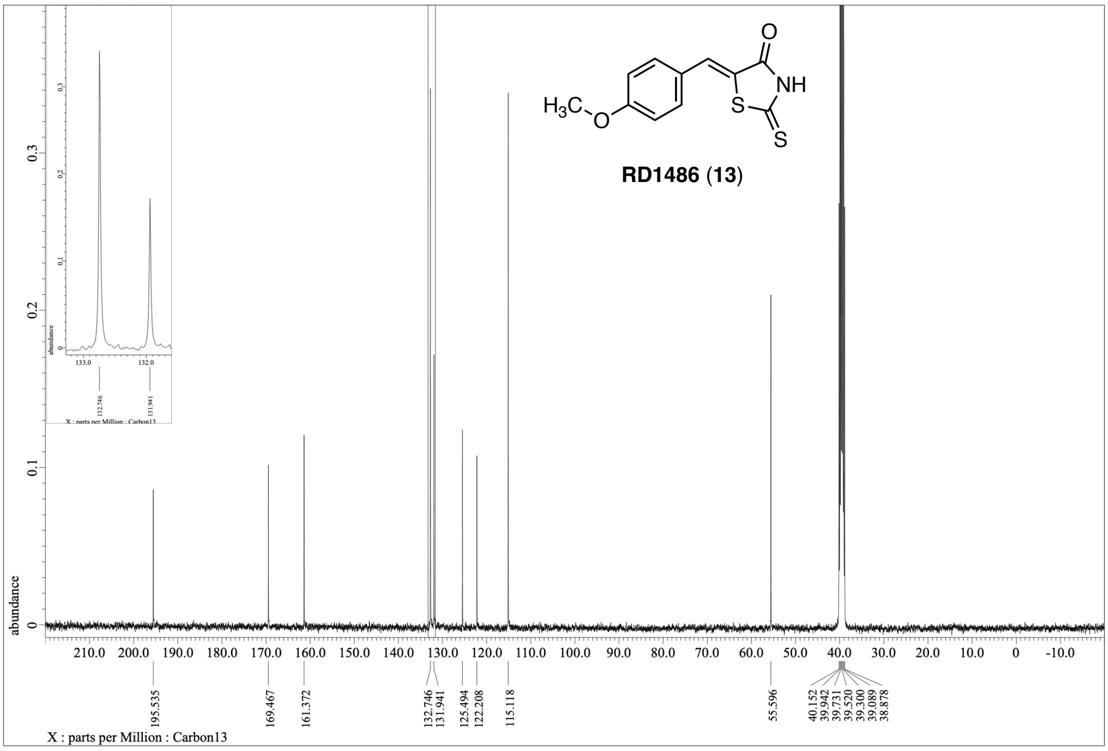

**Table S1.** Summary of $IC_{50}$ values for the indicated inhibitors calculated from the dose–response curves in Figures 2B–E,G–O (DYRK1A), 5B–N and S10C (DYRK1B).

| | $IC_{50}$ (μM) | | | |
|---|---|---|---|---|
| | DYRK1A | | DYRK1B | |
| Compound | – | Transient heating | – | Transient heating |
| FINDY (**1**) | > 32 | 11 | > 32 | 7.9 |
| *dp*-FINDY (**2**) | > 32 | 2.6 | > 32 | 2.5 |
| RD0447 (**3**) | > 32 | > 32 | > 32 | > 32 |
| RD1408 (**4**) | > 32 | > 32 | > 32 | 14 |
| RD0392 (**5**) | 6.1 | 4.8 | 3.6 | 3.5 |
| harmine (**6**) | 2.7 | 2.6 | 1.1 | 4.8 |
| staurosporine (**7**) | 0.40 | 0.42 | 0.097 | 0.096 |
| AZ191 (**8**) | 1.3 | 1.1 | 0.13 | 0.63 |
| GSK-626616 (**9**) | 0.60 | 0.90 | 0.49 | 3.2 |
| leucettine L41 (**10**) | 1.0 | 0.43 | 0.77 | 3.2 |
| RD0448 (**11**) | > 64 | 38 | 52 | 16 |
| RD0562 (**12**) | 31 | 17 | 23 | 23 |
| RD1486 (**13**) | 14 | 15 | 9.7 | 17 |

The $IC_{50}$ value represents the concentration of each compound that inhibits the kinase activity by 50%. Chemical structures of the compounds are shown in Figure 1B.

**Table S2.** Summary of $IC_{50}$ values for the indicated inhibitors calculated from the pre-incubation titration curves (DYRK1A) in Figure 4A–C and 4F,G.

| pre-incubation | $IC_{50}$ (μM) | | | | |
|---|---|---|---|---|---|
| time (hr) | FINDY (**1**) | *dp*-FINDY (**2**) | RD0392 (**5**) | leucettine L41 (**10**) | RD0448 (**11**) |
| 0 | – | – | – | 0.74 | – |
| 1 | > 32 | > 32 | 3.1 | – | > 64 |
| 2 | > 32 | > 32 | 4.0 | 0.63 | > 64 |
| 4 | 18 | > 32 | 2.9 | 0.52 | 46 |
| 6 | – | – | – | 0.52 | – |
| 8 | 12 | > 32 | 3.6 | 0.51 | 39 |
| 10 | – | – | – | 0.54 | – |
| 12 | 14 | > 32 | 3.5 | 0.45 | 37 |
| 16 | 15 | > 32 | 3.7 | – | 31 |
| 22 | 14 | 30 | 3.5 | – | 32 |
| 28 | 13 | 29 | 4.0 | – | 23 |
| 48 | 13 | 15 | 5.2 | – | 18 |

The $IC_{50}$ value represents the concentration of each compound that inhibits the kinase activity by 50%.

**Table S3.** Summary of $IC_{50}$ values for the indicated inhibitors calculated from the pre-incubation titration curves (DYRK1B) in Figure 5J–N.

| pre-incubation | $IC_{50}$ (μM) | | | | |
|---|---|---|---|---|---|
| time (min) | FINDY (**1**) | *dp*-FINDY (**2**) | RD0392 (**5**) | leucettine L41 (**10**) | RD0448 (**11**) |
| 0 | > 32 | > 32 | 2.4 | 0.53 | > 64 |
| 5 | 30 | – | – | – | – |
| 10 | 13 | – | – | – | – |
| 15 | 8.4 | – | – | – | – |
| 20 | 6.0 | 13 | 3.7 | 0.60 | 11 |
| 25 | 5.9 | – | – | – | – |
| 30 | 4.5 | – | – | – | – |
| 40 | – | 5.7 | 3.6 | 0.61 | 6.7 |
| 60 | – | 2.0 | 4.3 | 1.2 | 5.9 |
| 80 | – | 1.1 | – | – | 4.9 |
| 100 | – | 0.87 | – | – | 2.9 |
| 120 | – | 0.39 | – | – | 3.4 |

The $IC_{50}$ value represents the concentration of each compound that inhibits the kinase activity by 50%.

**Table S4.** Summary of $IC_{50}$ values for the indicated inhibitors calculated from the dose–response curves in Figures S4 (DYRK1A) and S10B (DYRK1B).

| | | $IC_{50}$ (μM) | | | |
|---|---|---|---|---|---|
| | | DYRK1A | | DYRK1B | |
| Compound | Structure | – | Transient heating | – | Transient heating |
| *mp*-FINDY | 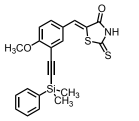 | > 32 | 4.5 | 30 | 2.1 |
| *tp*-FINDY | 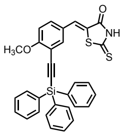 | > 32 | 29 | > 32 | 5.8 |
| *ts*-FINDY | 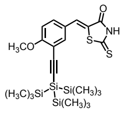 | > 32 | 9.9 | > 32 | 4.6 |
| *ms*-FINDY | 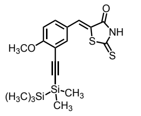 | > 32 | 2.6 | 28 | 0.8 |
| *et*-FINDY | 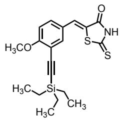 | > 32 | 1.4 | 14 | 0.90 |
| *tb*-FINDY | 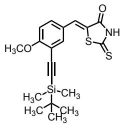 | > 32 | 1.8 | 17 | 1.5 |
| *ip*-FINDY | 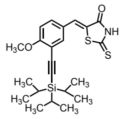 | > 32 | 4.7 | 21 | 1.2 |
| dFINDY | 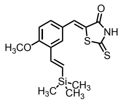 | > 32 | 4.9 | 20 | 1.9 |
| *dstb*-dFINDY | 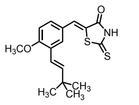 | > 32 | > 32 | > 32 | 7.4 |
| *dscy*-dFINDY | 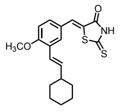 | > 32 | 4.1 | 27 | 2.5 |

The $IC_{50}$ value represents the concentration of each compound that inhibits the kinase activity by 50%. Chemical structures of the compounds are shown.

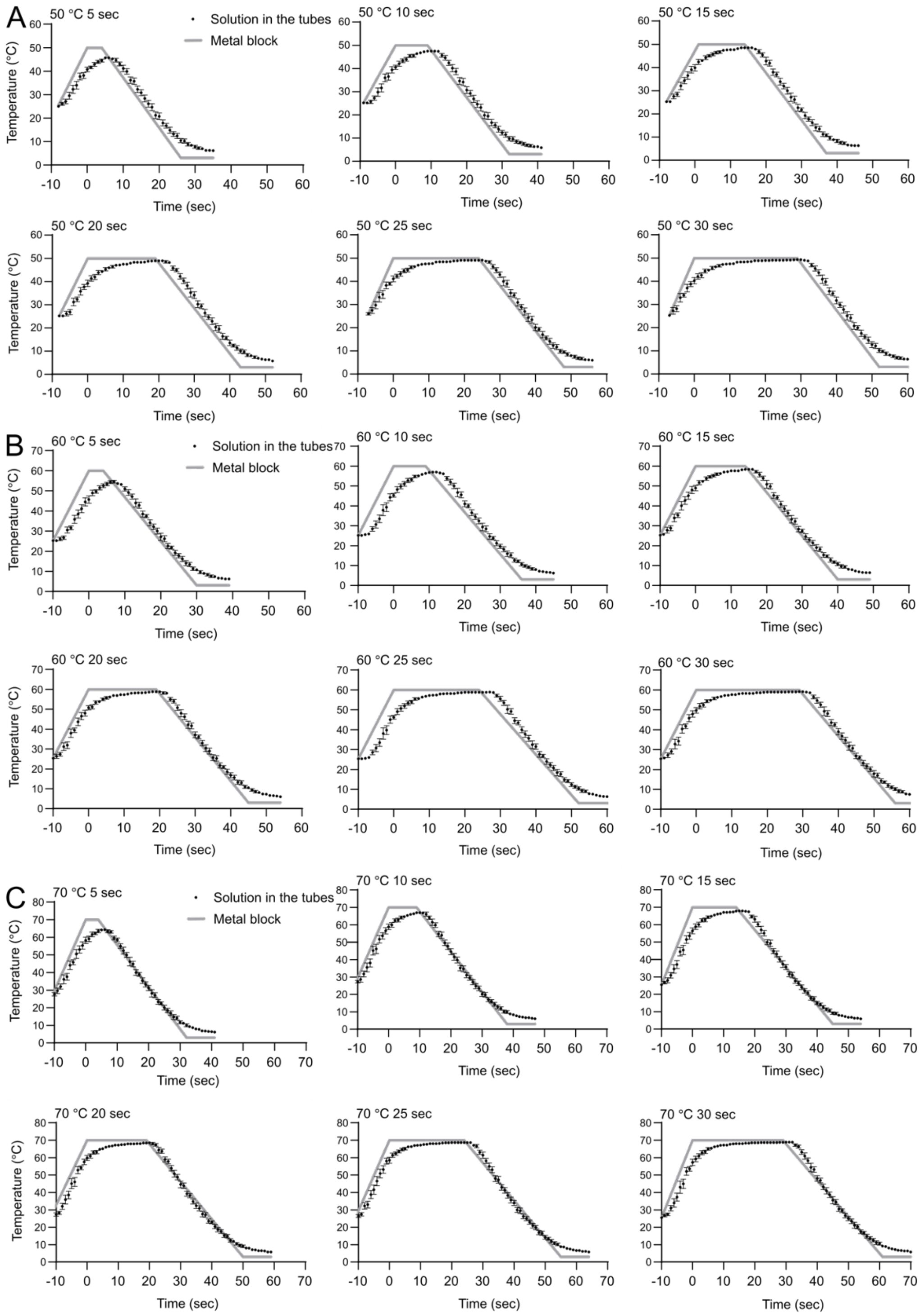


**Figure S1.** Actual temperature change of a solution in the thermal cycler.

(A–C) The temperatures of the solutions (15 µL) in a strip of eight PCR tubes in vertical column no. 7 of the thermal cycler were measured simultaneously. The heating condition was set to 50 ℃ (A), 60 ℃ (B), and 70 ℃ (C) for 5 sec, 10 sec, 15 sec, 20 sec, 25 sec, and 30 sec. The heating and cooling rates were set to the maximum of the thermal cycler. The data are presented as mean ± S.D. of the eight PCR tubes. The gray solid line indicates the temperature of the metal block, which was displayed on the thermal cycler.

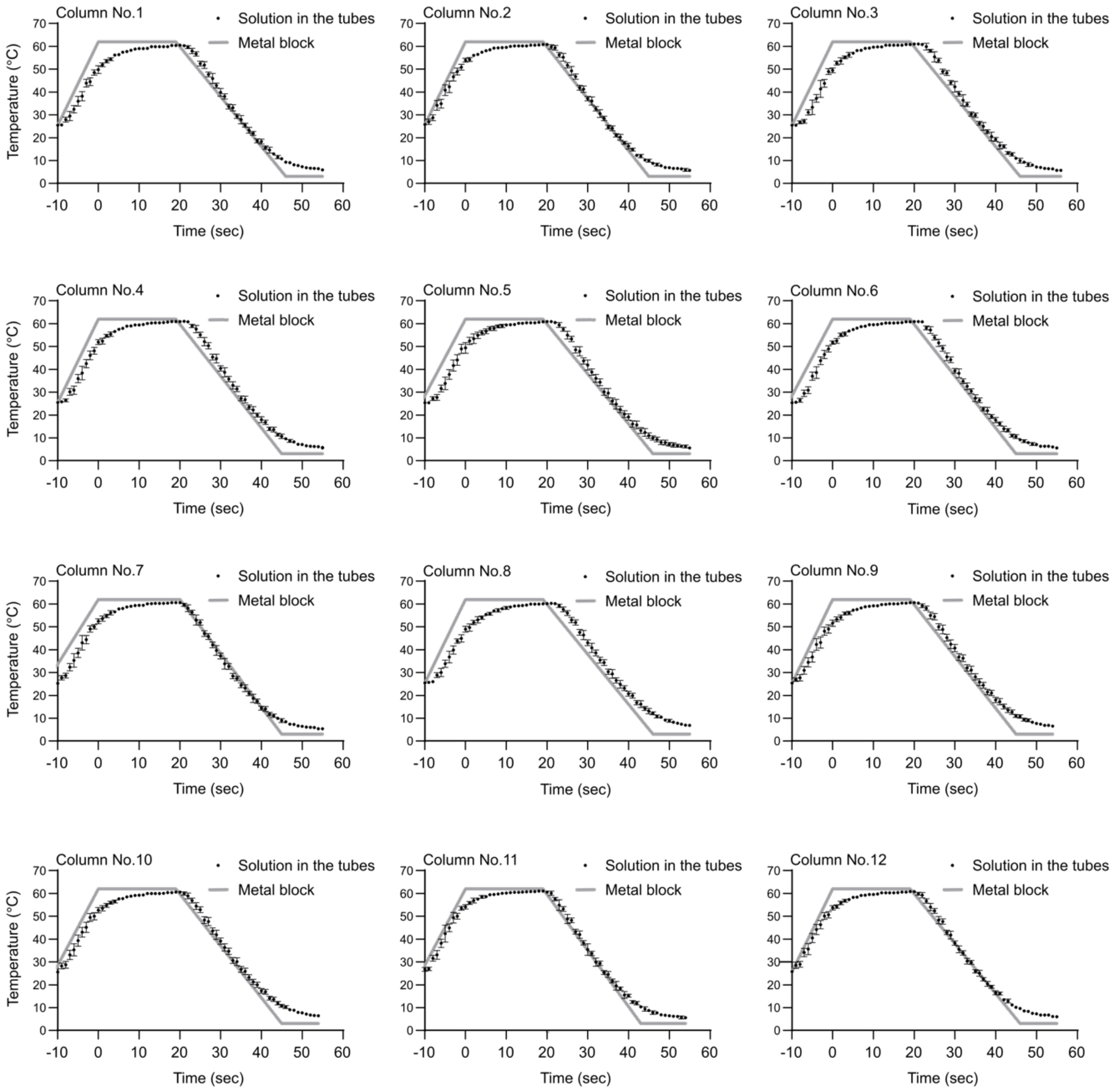


**Figure S2.** Validation of the stability and reproducibility of the temperature change in the 96 wells of the thermal cycler.

The temperatures of the solutions (15 μL) in a strip of eight PCR tubes in vertical columns no. 1–12 of the thermal cycler were measured simultaneously. The heating condition was set to 62 °C for 20 sec. The heating and cooling rates were set to the maximum of the thermal cycler at measured rates of 4.9 $°C \cdot sec^{-1}$ and 4.7 $°C \cdot sec^{-1}$, respectively. The data are presented as mean ± S.D. of the eight PCR tubes. The gray solid line indicates the temperature of the metal block, which was displayed on the thermal cycler.

A

| | 1 | 2 | 3 | 4 | 5 | 6 | 7 | 8 | 9 | 10 | 11 | 12 |
|---|---|---|---|---|---|---|---|---|---|---|---|---|
| A | 2397016 | 467456 | 567168 | 473672 | 372952 | 458848 | 323352 | 403320 | 354408 | 405864 | 516608 | 2247936 |
| B | 2200968 | 570224 | 591184 | 609720 | 494968 | 531216 | 377456 | 455576 | 468768 | 505704 | 527728 | 2054128 |
| C | 2149080 | 538960 | 593080 | 611232 | 512096 | 565720 | 330648 | 408432 | 416936 | 446760 | 628168 | 2058672 |
| D | 2279392 | 588800 | 456728 | 539176 | 559176 | 536832 | 359864 | 447136 | 399056 | 413840 | 432800 | 2040448 |
| E | 2316832 | 531208 | 525856 | 526216 | 624136 | 592296 | 463832 | 471856 | 384896 | 451960 | 517376 | 1937728 |
| F | 2271880 | 529032 | 516200 | 482152 | 437080 | 495352 | 499392 | 481872 | 451144 | 379368 | 565472 | 2005608 |
| G | 2225848 | 577680 | 515840 | 508800 | 536648 | 516800 | 499088 | 552064 | 513816 | 543248 | 581824 | 1975104 |
| H | 2185128 | 446984 | 453712 | 444384 | 478712 | 479152 | 472216 | 519096 | 451808 | 507336 | 560272 | 1956048 |

B

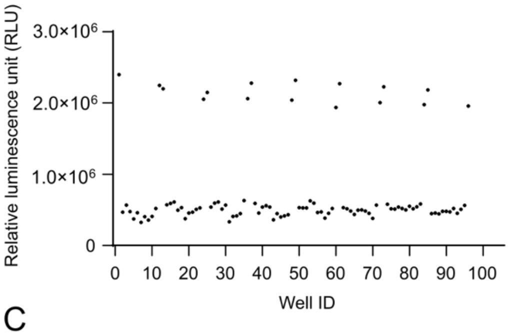


C

| | 1 | 2 | 3 | 4 | 5 | 6 | 7 | 8 | 9 | 10 | 11 | 12 |
|---|---|---|---|---|---|---|---|---|---|---|---|---|
| A | 11753824 | 1626520 | 1836800 | 1877464 | 1913992 | 2101312 | 1848392 | 1985800 | 1808584 | 1958728 | 1943672 | 11339464 |
| B | 11354640 | 1797392 | 1730240 | 1963328 | 1807120 | 1981032 | 1903632 | 1881016 | 1852560 | 1786584 | 1903872 | 11416704 |
| C | 11377944 | 1945120 | 1890088 | 1909672 | 1897080 | 2031264 | 1778528 | 1871680 | 1828840 | 1901776 | 2057232 | 10819984 |
| D | 11211896 | 2212592 | 1839328 | 1721016 | 1786296 | 2036952 | 1890480 | 1789512 | 1946384 | 1741888 | 1824896 | 10887256 |
| E | 11553824 | 1724744 | 1961960 | 1926280 | 1924096 | 2021272 | 1945320 | 1889584 | 1779600 | 1869408 | 1879752 | 9762888 |
| F | 11128560 | 1843824 | 1743432 | 1865464 | 1919432 | 1930392 | 1825008 | 1807992 | 1748224 | 1763496 | 1928288 | 10383120 |
| G | 11312992 | 1846448 | 1896752 | 1897304 | 1991416 | 2026320 | 1859336 | 1858304 | 1801896 | 1932856 | 1987688 | 10174968 |
| H | 11093344 | 1754008 | 1782224 | 1866952 | 1881800 | 2233688 | 1701176 | 1804880 | 1763864 | 1803440 | 2110448 | 10525128 |

D

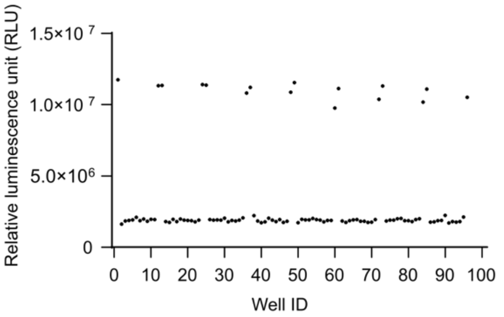


**Figure S3.** Robustness of the kinase assay with the temperature vaulting method.

The DYRK1A protein was transiently heated at 62 °C for 20 sec (A and B) or not (C and D) prior to the kinase reaction in a 96-well plate. A heat map representation shows the actual luminescence values (raw data) from the kinase assay with Kinase-Glo Luminescent Kinase Assay kit (A and C). Scatter plots of the actual luminescence values (raw data) from the kinase assay in 96-well format are shown (B and D). The horizontal axis indicates the well position in the 96-well format. The wells at position A-1 to H-1 and A-12 to H-12 indicate those in the kinase assay without the DYRK1A protein. The other wells included the DYRK1A protein in the kinase assay.

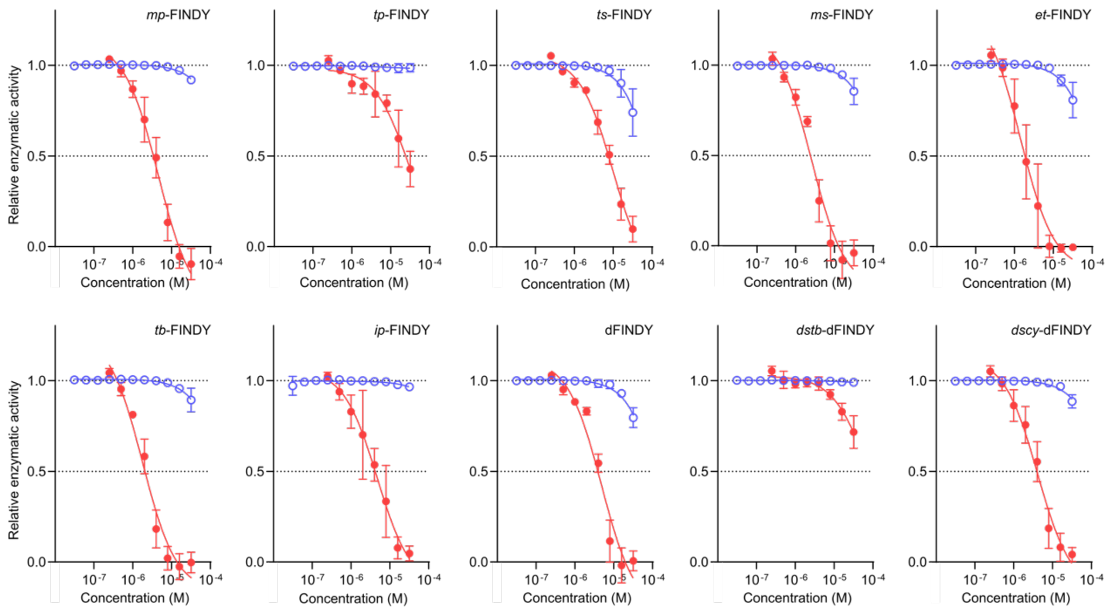


**Figure S4.** Inhibitory potency of the FINDY derivatives in the temperature vaulting method. The relative kinase activities of the DYRK1A protein in the presence of the FINDY derivatives, which were previously reported.[1] The red closed and blue open circles indicate the kinase activities with and without transient heating at 62 °C for 20 sec prior to the kinase reaction, respectively. Representative dose–response curves with Hill slopes are shown. The data are presented as mean ± S.D. of three independent experiments measured in duplicate. The $IC_{50}$ values are summarized in Table S4.

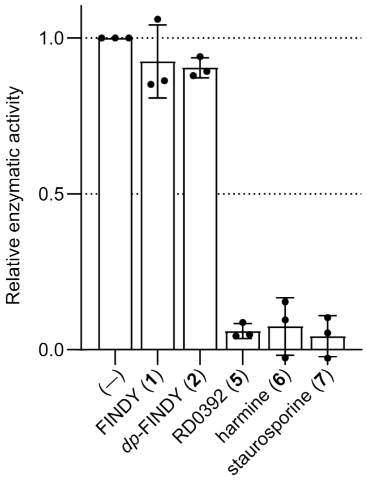


**Figure S5.** FINDY (**1**) and *dp*-FINDY (**2**) did not inhibit the DYRK1A protein that was once denatured and then refolded.

The DYRK1A protein transiently heated at 62 °C for 20 sec without any inhibitor was subjected to the kinase reaction in the presence of the indicated inhibitors (32 μM). The data are presented as mean ± S.D. of three independent experiments measured in duplicate.

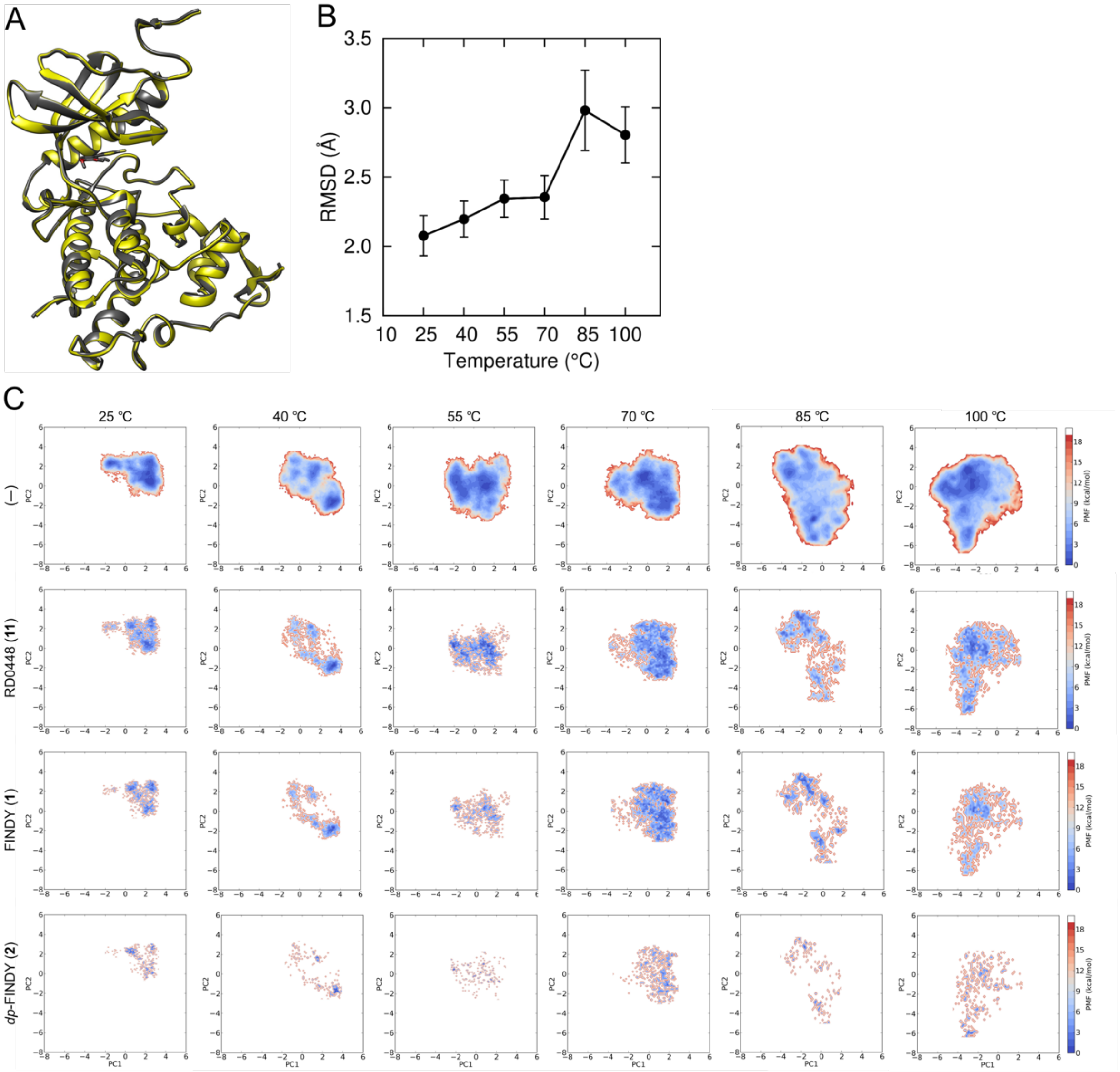


**Figure S6.** Analysis of the DYRK1A kinase domain complexed with the inhibitors.

(A) Overlay of the ribbon diagrams of crystal structures of DYRK1A kinase domain complexed with RD0392 (**5**) (PDB entry 7FHS; gray) and RD0448 (**11**) (PDB entry 7FHT; yellow).

(B) Effect of temperature on the protein backbone root-mean-square deviation (RMSD). The RMSD calculation was performed on the backbone from residues 144 to 481 (kinase domain) of DYRK1A at simulation temperature in the range of 25 to 100 °C. Values are averaged for the 2-µsec trajectory of four independent simulations.

(C) Top row columns show conformational distribution for DYRK1A kinase domain simulated at the indicated temperatures in PCA space. The color indicates potential of mean force (PMF). Blue and red spots indicate stable and unstable conformations, respectively. The bottom three columns show the conformational distribution for the simulated DYRK1A kinase domain that does not exhibit collision with the indicated inhibitor.

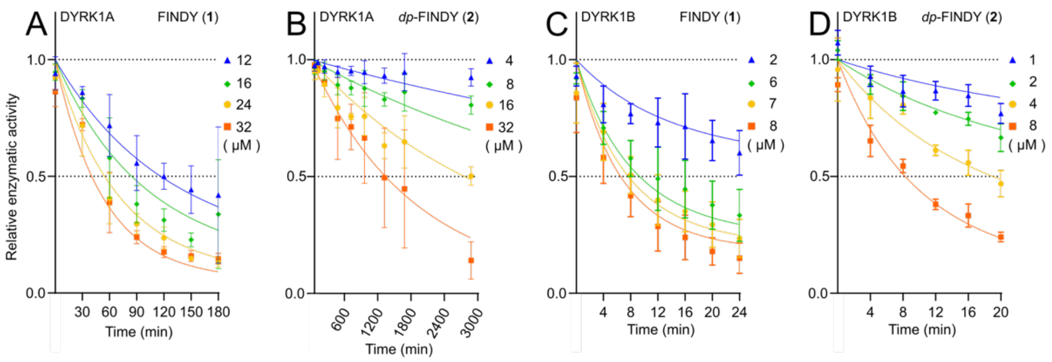


**Figure S7.** Inhibition of the DYRK1A/1B proteins by either FINDY (**1**) or *dp*-FINDY (**2**) under pre-equilibrium conditions.

The data in Figures 4A (DYRK1A/FINDY (**1**)), 5J (DYRK1B/FINDY (**1**)), and 5K (DYRK1B/*dp*-FINDY (**2**)) were globally fitted to eq 15,[4,5] but the resulting fit showed low accuracy. Therefore, we modified the inhibitor concentration range and pre-incubation time interval as shown in S7A (DYRK1A/FINDY (**1**)), S7C (DYRK1B/FINDY (**1**)), and S7D (DYRK1B/*dp*-FINDY (**2**)), re-collected and analyzed the data. The data are presented as mean ± S.D. of three independent experiments measured in duplicate. The data in Figure 4B (DYRK1A/*dp*-FINDY (**2**)) was globally fitted to eq 15 (S7B). The solid lines represent the best fit using eq 15. The dissociation constants ($K_D$), the association rate constants ($k_{on}$), the dissociation rate constants ($k_{off}$), the residence times, and the correlation coefficients of the global fitting ($R^2$) obtained from these data are presented in Table 1.

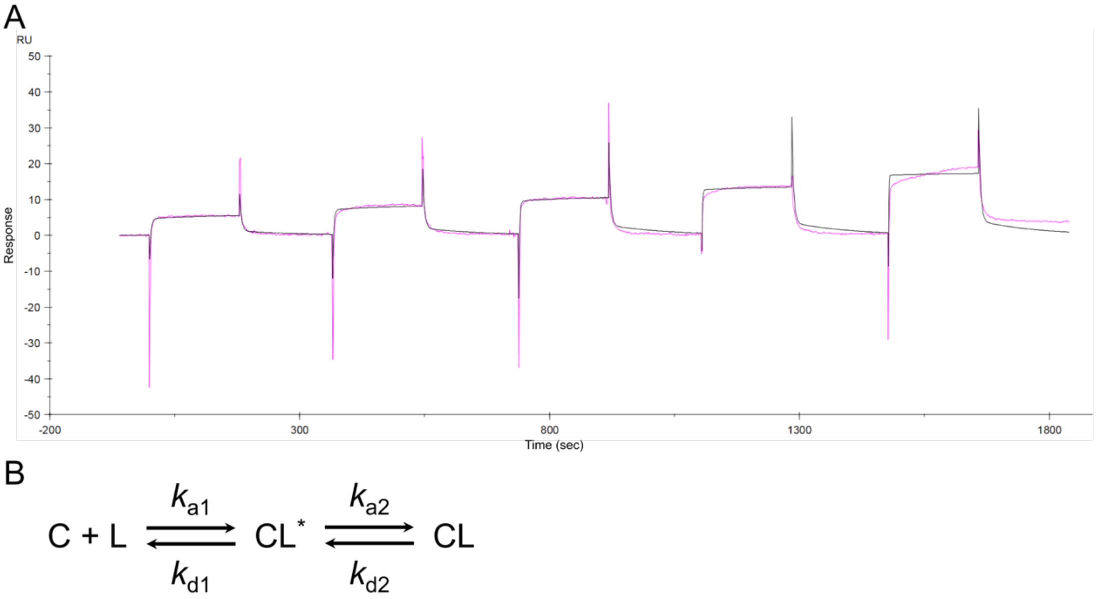


**Figure S8.** The association/dissociation kinetics of RD0392 (**5**) for the DYRK1A protein.
(A) Single-cycle kinetics analysis of surface plasmon resonance sensorgram for binding of the DYRK1A protein with RD0392 (**5**). Solutions of RD0392 (**5**) at concentrations of 50, 100, 200, 400, and 800 nM were injected over the immobilized DYRK1A protein (association phase), and then the dissociation of the complex was monitored. The black line represents the best fit using the two-state binding model.
(B) Two-state binding model. $k_{a1}$, $k_{a2}$, $k_{d1}$, and $k_{d2}$ are the rate constants for the indicated processes. For binding between the DYRK1A protein and RD0392 (**5**), $k_{a1}$ ($k_{on}$) is $2.0 \times 10^6$ $M^{-1}\cdot sec^{-1}$, $k_{d1}$ is 0.29 $sec^{-1}$, $k_{a2}$ is $1.0 \times 10^{-3}$ $sec^{-1}$, and $k_{d2}$ is $8.4 \times 10^{-3}$ $sec^{-1}$. $k_{off}$ is 0.26 $sec^{-1}$. $K_D$ is $1.3 \times 10^{-7}$ M. The data is the representative of three independent experiments.

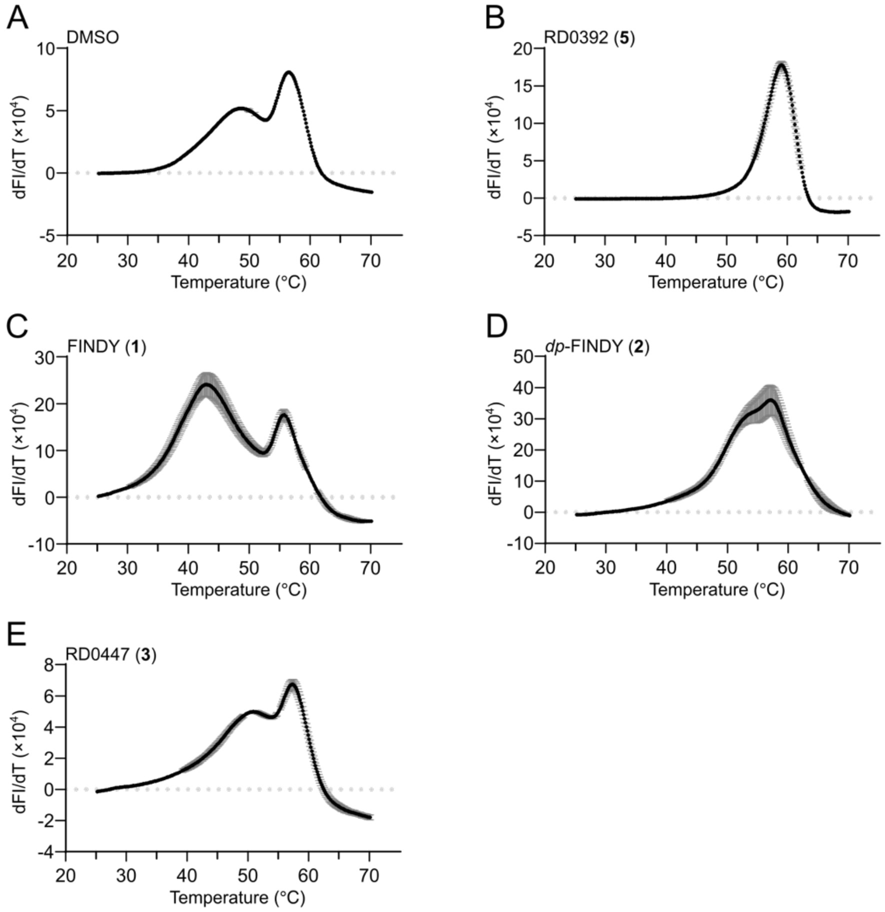


**Figure S9.** Thermal stability of the DYRK1A kinase domain in the presence of the inhibitors. Thermal stability analyses of the DYRK1A kinase domain in the presence of the solvent DMSO (A), RD0392 (**5**) (B), FINDY (**1**) (C), *dp*-FINDY (**2**) (D), and RD0447 (**3**) (E). The differences in the fluorescence intensity at each temperature (dFI/dT) are plotted against temperature. Peak temperatures in dFI/dT curves represent melting temperature ($T_m$). The data are presented as mean ± S.D. of three independent experiments.

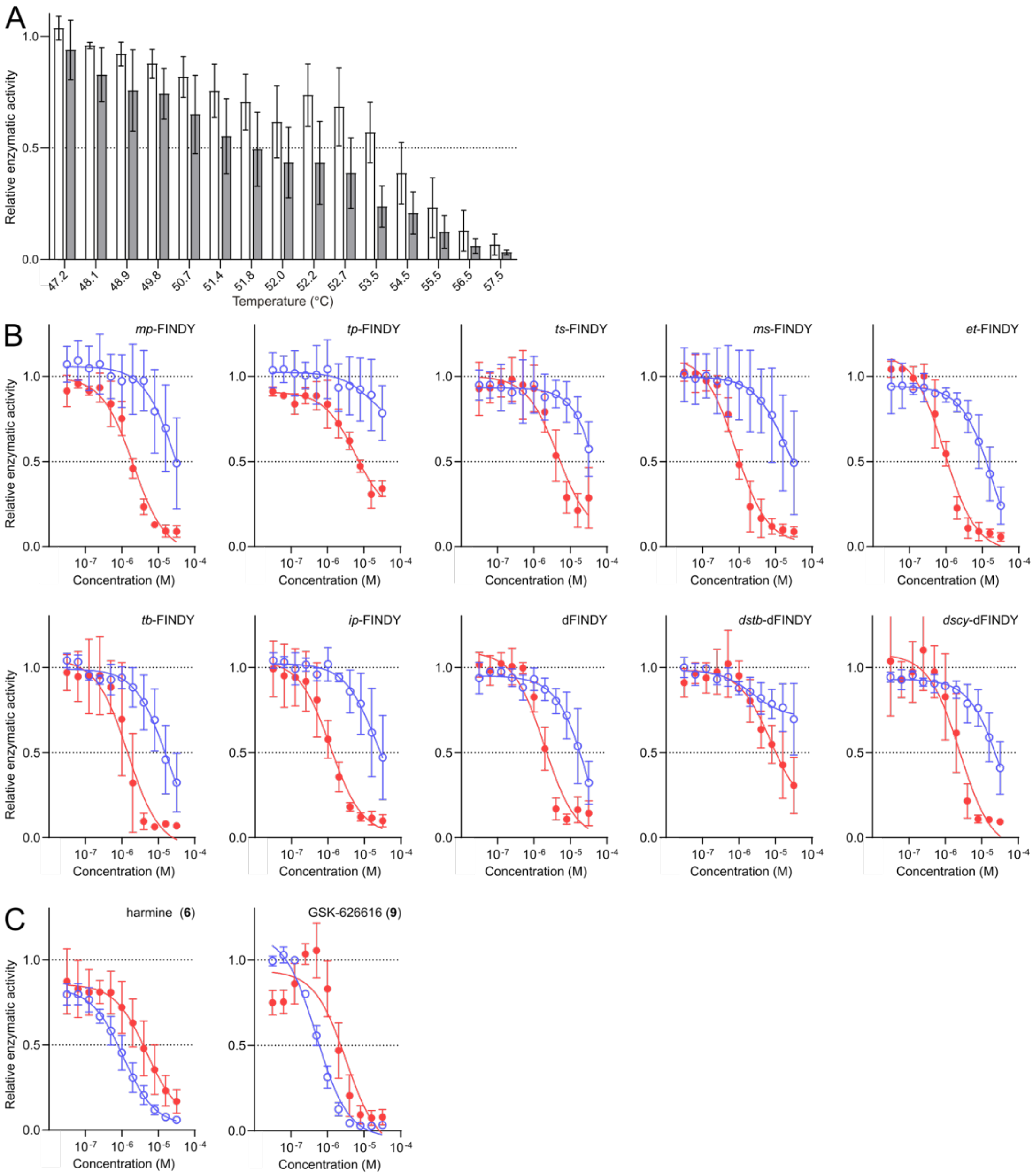


**Figure S10.** The transient-heating-dependent inhibition of the DYRK1B protein.

(A) Determination of the optimum heating temperature. The DYRK1B protein was transiently heated at the indicated temperatures for 20 sec in the presence and absence of FINDY (**1**) at 8.0 µM (closed and open bars, respectively). The kinase activities relative to that at 37 °C in the absence of FINDY (**1**) are displayed on the bar graphs. The data are presented as mean ± S.D. of three independent experiments measured in duplicate.

(B) Inhibitory potency of the FINDY derivatives in the temperature vaulting method. The relative kinase activities of the DYRK1B protein in the presence of *mp*-FINDY, *tp*-FINDY, *ts*-FINDY, *ms*-FINDY, *et*-FINDY, *tb*-FINDY, *ip*-FINDY, dFINDY, *dstb*-dFINDY, and *dscy*-dFINDY. The red closed and blue open circles indicate the kinase activities with and without transient heating prior to the kinase reaction, respectively. Representative dose–response curves with Hill slopes are shown. The data are presented as mean ± S.D. of three independent experiments measured in duplicate. The $IC_{50}$ values are summarized in Table S4.

(C) The relative kinase activities of the DYRK1B protein in the presence of harmine (**6**) and GSK-626616 (**9**). The red closed and blue open circles indicate the kinase activities with and without transient heating prior to the kinase reaction, respectively. Representative dose–response curves with Hill slopes are shown. The data are presented as mean ± S.D. of three independent experiments measured in duplicate. The $IC_{50}$ values are summarized in Table S1.
